# An extended structure of the intracellular domain of the Torpedo nicotinic acetylcholine receptor and its proposed interactions with rapsyn

**DOI:** 10.64898/2025.12.03.692134

**Authors:** Camille M. Hénault, Moustafa Habes, Christian J.G. Tessier, John Baenziger

## Abstract

To gain insight into the interactions between rapsyn and the nAChR that induce clustering at the post-synaptic membrane, we refined a cryo-EM dataset using an intracellular domain focused strategy to obtain a 3.0 Å map with the most extensive density yet for the intracellular domain of the *Torpedo* nAChR. The improved map allowed us to extend the structure beyond the MX α-helix and prior to the MA α-helix of the intracellular domain. The new structure defines a sharp N-terminal boundary of each MA α-helix to place agrin-dependent phosphorylated tyrosines unambiguously within the flexible regions of the MX-MA loops. Two distinct conformations of the α_δ_ M4 α-helix were also resolved, indicating that M4 conformational heterogeneity reflects intrinsic flexibility rather than a change in gating state. The new structural constraints defined for the MX-MA loop were then used to evaluate AlphaFold3-predicted full-length models of the nAChR, rapsyn, and various rapsyn-nAChR complexes, identifying a consistent, asymmetric 3:1 binding architecture where each rapsyn is always sandwiched between the MX-MA loops from two subunits and where each phospho-tyrosine is lodged in a cationic pocket formed by conserved residues implicated in congenital myasthenic syndromes. The defined architecture fits published cryo-ET maps of *Torpedo* post-synaptic membranes and explains how both phosphorylated tyrosines and myasthenic syndrome-causing rapsyn mutations modulate receptor clustering.

## Introduction

Rapid communication at the neuromuscular junction depends upon the clustering of nicotinic acetylcholine receptors (nAChRs) in regions of the postsynaptic muscle membrane that align with active zones in the presynaptic motor neuron. Clustering at the developing neuromuscular junction is initiated by the release of the proteoglycan, agrin, from the presynaptic nerve terminus (Froehner, 1993; Glass & Yancopoulos, 1997; Ferns, 2021). The released agrin binds to the post-synaptic low-density lipoprotein-related protein 4 (LRP4) receptor, which activates its co-receptor, the muscle specific protein kinase (MuSK), to initiate intracellular signaling cascades that ultimately lead to phosphorylation of the nAChR (Kim et al., 2008; Walker et al., 2021; Xie et al., 2023). nAChR phosphorylation promotes interactions between the nAChR and the receptor associated protein of the synapse, rapsyn, with the bound rapsyn in turn interacting with other rapsyn-nAChR complexes and/or with other cytoskeleton proteins to facilitate nAChR clustering at the top of the post-synaptic junctional folds (Prömer et al., 2023).

Building on cryo-electron microscopy (cryo-EM) reconstructions from helical tubes of the *Torpedo* nAChR and cryo-electron tomography (cryo-ET) maps of *Torpedo* post-synaptic membranes (Unwin, 2024; Zuber & Unwin, 2013), recent structures of both the muscle-type nAChR (Rahman et al., 2022; Zarkadas et al., 2022) from *Torpedo* and the mammalian adult/fetal muscle nAChRs (H. Li et al., 2024) provide a framework for understanding the nature of rapsyn-nAChR interactions. Muscle nAChRs are formed from five homologous subunits arranged in an α-β-δ-α-γ/ε clockwise fashion, with the mammalian fetal γ subunit replaced by the adult ε subunit during later muscle development. Each of the subunits includes an extracellular domain (ECD) formed predominantly from 10 β-strands, β1-β10, a transmembrane domain (TMD) formed from four transmembrane α-helices, M1-M4, and an intracellular domain (ICD) extending between M3 and M4 that consists of a short MX α-helix, located at the interface between the lipid bilayer and cytoplasm, followed by an unstructured region and then a long MA α-helix that is contiguous with M4. The binding of acetylcholine (ACh) occurs in the ECD at the interfaces between the two principal α subunits and their complimentary δ and γ/ε subunits, while channel gating occurs in the TMD and is driven by movements of the five pore-lining M2 transmembrane α-helices. Although not fully resolved in any nAChR structure solved to date, the five MA α-helices and the unstructured regions of the ICD are thought to facilitate interactions with rapsyn to promote nAChR clustering.

In contrast to the nAChR, the structure of rapsyn remains poorly defined (Ponting & Phillips, 1996; Ramarao et al., 2001). Sequence analysis originally predicted an N-terminal myristylation site, seven tetracopeptide repeats (TPRs), a coiled-coil domain, and a RING-H2 domain (Frail et al., 1988), with functional deletions studies suggesting a role for each in nAChR clustering. The N-terminal myristylation site anchors rapsyn to the plasma membrane (Phillips et al., 1991) while the TPR motifs mediate clustering through both rapsyn-rapsyn self-association and rapsyn interactions with other post-synaptic proteins (Colledge & Froehner, 1998). The coiled-coil domain also facilitates nAChR-rapsyn interactions (Ramarao et al., 2001) while the E3 ligase activity of the RING-H2 domain is responsible for nAChR neddylation, which slows nAChR turnover rates (Xing et al., 2019).

The structural basis for rapsyn-nAChR interactions also remain poorly understood with even the stoichiometry of the interactions still debated. Although early biochemical studies suggested a 1:1 molar stoichiometry for rapsyn binding to the nAChR (Burden et al., 1983; LaRochelle & Froehner, 1986), increasing evidence suggests higher, yet variable interaction stoichiometries of up to three rapsyns per nAChR (Moransard et al., 2003; Gervásio et al., 2007; Brockhausen et al., 2008; Ferns, 2021; Martínez-Martínez et al., 2007; Zuber & Unwin, 2013). It is not even clear which nAChR subunits interact directly with rapsyn. Some studies suggest binding to the α, β, and ε/γ (Lee et al., 2009) or to the β, δ, and ε/γ subunits (Huebsch & Maimone, 2003). Other studies suggest binding to all four subunits (Maimone & Merlie, 1993) or to the interfaces between multiple subunits (Ferns, 2021). Even the regions of each ICD that bind rapsyn remain unclear (Huebsch & Maimone, 2003), with some suggesting that the MA α-helices are directly implicated (Maimone & Merlie, 1993; Huebsch & Maimone, 2003) while others suggest that the interactions are facilitated by nearby phosphorylated tyrosine residues (Borges et al., 2008).

Both the low solubility of rapsyn and its inherent propensity to self-aggregate and/or interact with other proteins continue to limit our structural understanding of both rapsyn and rapsyn-nAChR interactions, and thus our understanding of the mechanisms that underlie nAChR clustering at the neuromuscular junction. Our structural understanding of rapsyn-nAChR interactions and thus clustering is further limited by the incomplete structural characterization of the nAChR ICD (see, however, (Bondarenko et al., 2022) and by the inherent difficulties associated with structural studies of a complex that may exists with variable stoichiometries of interacting partners.

We recently recorded cryo-EM images of the *Torpedo* nAChR across a range of acetylcholine (ACh) concentrations, with combined datasets leading to reconstructions for un-, mono-, and di-liganded states (Thompson et al., 2025). Among these, the dataset recorded at 50 µM ACh yielded a map with relatively well-defined density for the intracellular domain (ICD). Here we refine this map further to resolve new structural elements between the MX and MA α-helices of each subunit that constrain the conformational space of an intrinsically disordered stretch of each ICD (Fig. 1). Building on these new constraints, we then predict full-length models of the nAChR, rapsyn, and the nAChR-rapsyn complex, the latter validated by comparison with our new ICD structures and published cryo-electron tomography (cryo-ET) maps of *Torpedo* postsynaptic membranes (Zuber & Unwin, 2013). Our new model visualizes a structure-based explanation for the architectural asymmetry of rapsyn binding to the nAChR that is observed in cryo-ET maps of *Torpedo* post-synaptic membranes. The predicted *Torpedo* rapsyn-nAChR complex also suggests that the agrin-mediated phosphorylated tyrosines residue in each of the β, γ, and δ subunit ICDs binds to a narrow pocket in rapsyn lined with positively charged residues that are implicated in the muscle disease, congenital myasthenic syndrome (CMS). In fact, the predicted binding mode mimics the mechanisms by which homologous TPR-motif proteins, such as Pex5, recognize their binding targets. Together, the new nAChR structure and the predicted models of rapsyn-nAChR interaction provide a starting framework for understanding both agrin-dependent clustering of the nAChR in post-synaptic membranes and how mutations in rapsyn lead to CMS.

**Figure 1.**
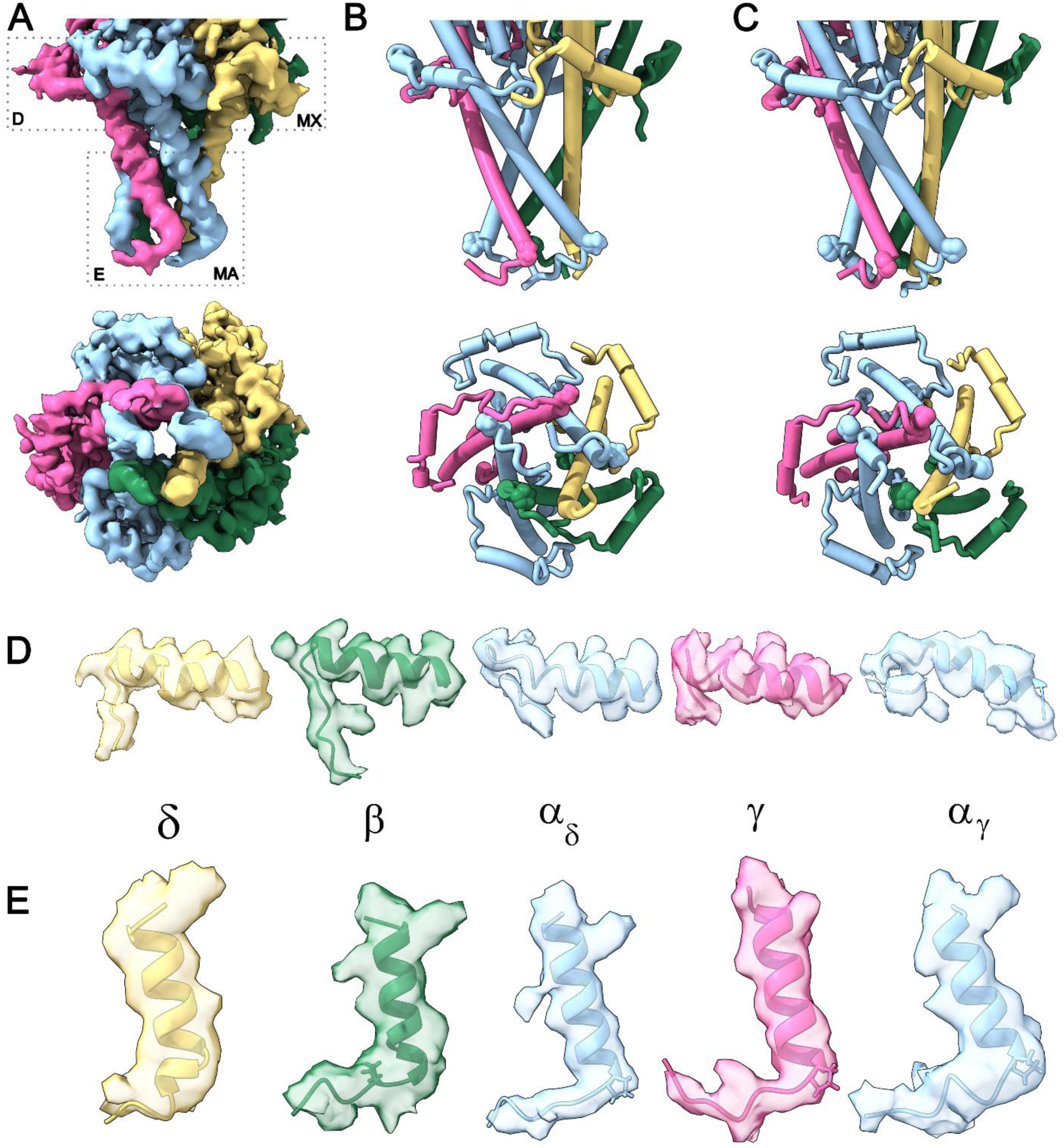
Enhanced structure of the nAChR ICD. **(A)** Side and bottom-up/cytoplasmic views of the enhanced ICD map highlighting the structures in the MX and MA regions. Subunits are colored α_δ_ and α_γ_, light blue; β, green); γ, pink; and δ, yellow. **(B)** Side and bottom-up/cytoplasmic views of the enhanced ICD model. **(C)** Side and bottom-up/cytoplasmic views of an averaged structure derived from ten AlphaFold3 predictions of the *Torpedo* nAChR alone (no phosphorylation or rapsyn) showing the close correlation between features in the predictions and those observed in our enhanced ICD structure. Zoomed in views of the MX **(D)** and MA **(E)** α-helices from each subunit fit into their respective cryo-EM densities.

## Results

### Extended ICD structure of the *Torpedo* nAChR

To extend our understanding of the structure of the nAChR ICD, we focused on a cryo-EM data set recorded at 50µM ACh from the *Torpedo* nAChR reconstituted into asolectin MSP2N2 nanodiscs (Fig. 1A-E). The original processing of the cryo-EM data yielded a 2.8-Å resolution map (Thompson et al., 2025) with more density for the ICD than observed in other maps of the *Torpedo* nAChR (Rahman et al., 2022; Zarkadas et al., 2022). To improve the coverage of the ICD further, we reprocessed the entire data set performing additional rounds of 3D-classifications using volume masks surrounding either the ICD alone or the ICD (Fig. S3) plus various regions of the TMD (Punjani et al., 2017; Punjani & Fleet, 2021). Refinement using a mask surrounding only the ICD ultimately led to a map at 3.0 Å resolution map (∼140,000 particles) with the most complete ICD density to date, referred to as the “ICD map” (Figs. 1, 2 & S1-S3).

**Figure 2.**
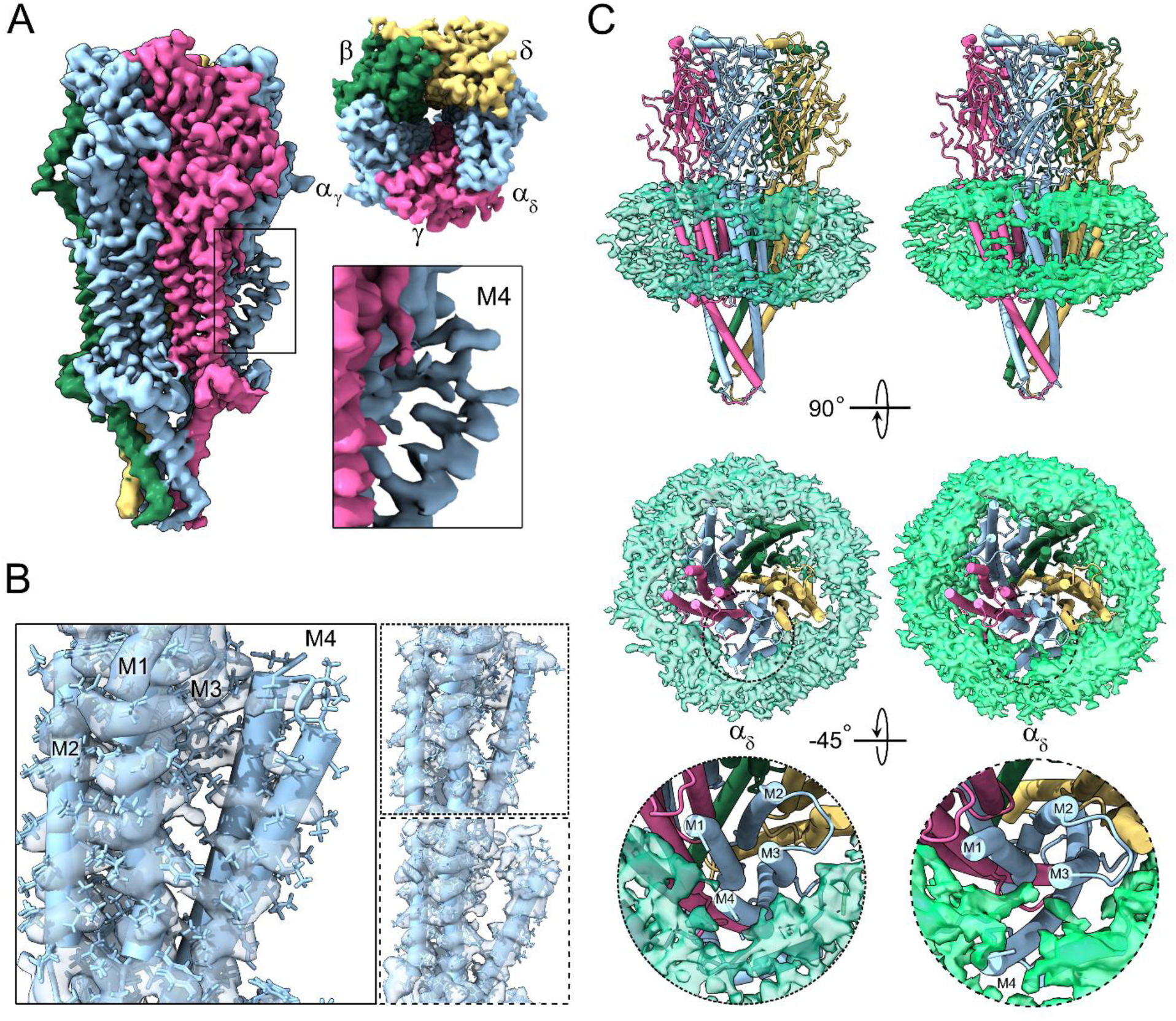
Cryo-EM maps and models of the *Torpedo* nAChR after refinement using a mask focused on the ICD. **(A)** Side and top views of the 3.04 Å ICD map. Individual subunits are colored α_γ_ and α_δ_, light blue; β, green; δ, yellow; and γ, pink. The box shows a zoomed in view of the α_δ_ M4 α-helix. **(B)** Zoomed in side view of the α_δ_ ICD map with the α_δ_M4-tight and α_δ_M4-tilt models overlaid (left). Insets to the right show the α_δ_M4-tight (short dashes) and α_δ_M4-tilt (long dashes) models and maps. **(C)** Side (top row) and top-down (middle row) views of the α_δ_M4-tight (left column) and α_δ_M4-tilt (right column) models fit into their nanodisc densities. The bottom row shows a zoomed in view of both models and their nanodisc maps looking straight down at the α_δ_M4 TMD to highlight both the relative position of the α_δ_M1 – α_δ_M4 α-helices and the position of the α_δ_M4 within the nanodisc.

The resulting “ICD structure”, which was generated by fitting the nicotine-bound structure (PDB:7QL5) into the unsharpened ICD map, adopts the typical pLGIC fold, with five subunits arranged in a clockwise α_γ_–β–δ–α_δ_–γ fashion (Figs. 1 and 2). As expected, the structure superimposes well onto a recently published ACh-bound *Torpedo* nAChR structure (PDB: 9E3E) (RMSDs values between 0.52 and 0.65 Å). Loops in the ECD that are involved in agonist-induced activation, the C- and F-loops, are capped around strong ACh density in both the α_γ_γ and α_δ_δ binding sites. Relative to the apo resting state, the pore-lining M2 α-helices are tilted/twisted away from the long axis of the central ion pore leading to a dilated L9’ gate. Although the nAChR is expected to adopt a desensitized conformation in the presence of 50 μM ACh, previous molecular dynamics simulations suggest that the ACh-bound structure exhibits a hydrated pore that conducts cations across the membrane (Thompson et al., 2025). It has been proposed that lipids lodged into the pore during sample vitrification trap the pore in a conformation along the trajectory between open and desensitized states (Zarkadas et al., 2022).

The ICD structure includes an additional 76 residues in the ICD compared to our reference nicotine bound structure (PDB:7QL5) and 45 residues relative to the more recently published ACh-bound structure (PDB: 9E3E) obtained from the same data set but prior to our refinement focused on the ICD (Figs 1 & S6). While the number of added residues is limited in the context of the total size of the ICD, the additional residues extend the structure of the polypeptide backbone C-terminal to each MX α-helix by a total of 21/18 total residues relative to the published nicotine-/ACh-bound structures. Moreover, the new map clearly defines the start/N terminus of each MA α-helix, as well as the path of the polypeptide backbone six or seven residues preceding MA in each subunit (Fig. 1). Of the added 76 residues, 37 located primarily in MA were modelled in a region of density where the side chain positions were interpreted with confidence (Figs. S4-S6). An additional 27 residues, mainly at the N-termini of the MA α-helices fall within an intermediate resolution where the backbone position is clear, but side-chain positions were less definitive. The remaining 12 residues (α_γ_K334 (MX), α_γ_T364, α_γ_P365, α_γ_L366, βP394, βV395, βT396, δD413, δQ414, γD405, γA407, γN408) located post-MX exhibit density that allowed definition of the path of the polypeptide backbone, with side-chain positions uncertain. We included the latter in our model only in so far as they help define the overall architecture of the ICD in this region.

C-terminal to each MX α-helix, each polypeptide backbone turns sharply downwards from the bilayer for between three to five residues ultimately projecting towards the sequence that precedes each MA α-helix (Fig. 1D). This downward turn is also seen in some subunits of other recently published structures (H. Li et al., 2025). After this downward projection, each MX – MA loop adopts a structurally disordered or flexible stretch, which is not seen in our cryo-EM maps or other cryo-EM maps published to date, before projecting parallel to the bilayer surface and then turning abruptly upwards to join each MA α-helix (Fig. 1E). The sharp turn upwards in the α, β, and γ subunits is facilitated by a proline residue (αPro370, βPro398, and γPro411) that defines the start of each MA α-helix (Fig. 1B). The δ ICD lacks an analogous proline residue with the path of the polypeptide backbone preceding MA tilted away from a membrane parallel direction so that it undergoes a shallower bend to join up with MA. Notably, the new ICD structure shows that the N and C termini of each disordered loop are oriented towards each other, which places constraints on the conformational space available to each MX-MA loop (Fig. 1). The defined N-termini of each MA α-helix also show unequivocally that the key agrin-dependent phosphorylated tyrosine residues, βTyr390, γTyr364, and δTyr393, are not part of an extended MA α-helical structure. This observation confirms that it is the loop between MX and MA, not an extended MA α-helix itself, in the β, γ, and δ subunits that facilitates agrin-mediated interactions between the nAChR and rapsyn.

The five MA α-helices together form an inverted cone-like structure with the base of the cone located near the bilayer surface and the apex of the cone projecting roughly 40 Å into the cytoplasm (Fig. 1A, B). At the apex of the cone-like structure, the N termini of the five MA α-helices are arranged around the long central axis of the nAChR overlapping like the blades in the iris of a camera. The N termini of the five MA α-helices form a regular pentagon lined by hydrophobic residues, αVal372, βLeu401, γIle413, and δLeu419. Distances across this pentagon are greater than 10 Å creating a gap that should be sufficient to pass hydrated cations. The flow of cations, however, is blocked by the α_γ_, δ, and α_δ_ subunit MX-MA loops leading into each MA α-helix, which pass directly under the apex of the inverted MA α-helix cone to fill and thus cap the opening between the MA α-helix N termini. In contrast, the β and γ subunit MX-MA loops leading into each MA α-helix project tangentially to the apex of the MA α-helix inverted cone such that they interact with the adjacent α_γ_, δ, and α_δ_ polypeptide chains, possibly to stabilize the cap structure. As discussed below, elements of this ICD organization are captured in predicted models of the nAChR-rapsyn complex where they facilitate interactions with rapsyn.

### Conformational variability in the M4 α-helix

During initial map refinement, we observed that the M4 α-helix region from the α_δ_ subunit (α_δ_M4) exhibits ambiguous density suggestive of two conformations (Figs 2A & S2-S3). To address this heterogeneity, we separated the particles into two main conformations, with further refinement yielding maps referred to as α_δ_M4-tight and α_δ_M4-tilt (Fig. 2B, C). The α_δ_M4-tight map at 3.2 Å resolution (∼50,000 particles) has the α_δ_M4 density packed against the densities from the adjacent α_δ_M1 and α_δ_M3 α-helices. The α_δ_M4-tilt map at 3.1 Å resolution (∼90,000 particles) has the α_δ_M4 density tilted away from α_δ_M1 and α_δ_M3, in a pose reminiscent of recently published α_δ_M4 maps and models (Rahman et al., 2022). We observed no correlation between the degree of α_δ_M4 tilt and the degree to which the ICD is structured and thus resolved in the cryo-EM maps. Similar conformational heterogeneity was not detected with any of the other M4 α-helices, suggesting that the M4 conformational variability is a α_δ_-specific phenomenon.

The similar numbers of particles contributing to the α_δ_M4-tight and α_δ_M4-tilt conformations (roughly 1:2, respectively) suggests that the two α_δ_M4 poses are not strict markers of a particular gating state, particularly given that both maps were derived from the same data set recorded at saturating ACh concentrations. This interpretation is supported by the observation that in both the α_δ_M4-tight and α_δ_M4-tilt reconstructions, the nAChR is positioned in the lipid nanodisc so that the α_δ_M4 α-helix is close to diffuse density that we attribute to the nanodisc scaffold protein, MSP2N2. In the α_δ_M4-tight map, this putative nanodisc density is in contact with the outer face of the α_δ_M4 α-helix such that all the observed density between the densities attributed to the polypeptide backbones of α_δ_M4 and α_δ_M1/α_δ_M3 is fully accounted for by side chains. In the α_δ_M4-tilt map, the α_δ_M4 density is tilted out towards the putative MSP2N2 density, with the density for the final few α_δ_M4 residues ambiguous. New density due to either lipid or MSP2N2 itself is also seen between the α_δ_M4 projection and both α_δ_M1 and α_δ_M3.

The two structures with distinct α_δ_M4 conformations, referred to as α_δ_M4-tight and α_δ_M4-tilt, were derived from the same dataset recorded in the presence of 50 μM Ach (Fig. 2). Not surprisingly, both structures correspond to the same functional di-liganded state with their C- and F-loops capped around bound ACh and their pore-lining M2 α-helices tilted/twisted away from their apo conformation leading to a dilated L9’ pore gate. This observation shows that the same functional state of the nAChR can exhibit different α_δ_M4 poses, at least when the nAChR is reconstituted into a lipid nanodisc. Our structures suggest that the two conformations are stabilized by distinct interactions between α_δ_M4 and the nanodisc scaffolding protein, MSP2N2. In the α_δ_M4-tight structure, α_δ_M4 is oriented close to the adjacent α_δ_M1 and α_δ_M3 α-helices so that the diffuse MSP2N2 and/or lipid density is peripheral to the α_δ_M4 C terminus. In the α_δ_M4-tilted reconstruction, the α_δ_M4 extends directly toward the MSP2N2 so that the scaffolding protein or lipid density is now positioned closer to α_δ_M1/α_δ_M3 (Fig. 2C).

Although early structures of the *Torpedo* nAChR suggested that α_δ_M4 adopts a distinct conformation in the desensitized state (Rahman et al., 2022), the α_δ_M4-tight and α_δ_M4-tilt conformations support an increasing body of structural data obtained from the nAChR and other pLGICs, most notably the prokaryotic pLGIC, ELIC, showing that there is intrinsic conformational flexibility of the outermost M4 transmembrane α-helix (Dalal et al., 2024). Interestingly, the outermost M4 α-helix of ELIC can adopt conformations that resemble both the α_δ_M4-tight and α_δ_M4-tilt conformations, with larger nanodiscs promoting the tilted M4 to also facilitate interactions between the M4 C-terminus and the nanodisc scaffolding protein (Dalal et al., 2024). Structures of the *Torpedo* nAChR with an increased M4 tilt have also been observed in cryo-EM images recorded from native tubular 2D crystals of *Torpedo* membranes, which exhibit a higher degree of intrinsic membrane curvature (Unwin, 2024). The intrinsic conformational flexibility of M4, which may underlie the exquisite lipid-sensitivity of the *Torpedo* nAChR pLGICs (daCosta et al., 2009, 2013), is likely an inherent property of some. The different α_δ_M4 tilt conformations observed in our data set likely reflect both this intrinsic flexibility and an extrinsic stabilization through interactions with the nanodisc scaffolds proteins, rather than a functional conformation-specific rearrangement.

### Predictive models in the context of the extended ICD structures

Although a substantial region of each MX-MA loop remains undefined in our new structures, the extended ICD density clearly resolves both the N- and C-terminal structural boundaries of each MX-MA loop, thereby constraining its conformational space. To visualize this space and how it might shape rapsyn-nAChR interactions, we used AlphaFold3 to predict full-length models of both the *Torpedo* nAChR, rapsyn, and various rapsyn-nAChR complexes (Figs. 3, 4 & S7). We generated over 200 models in total, with ∼85% of the models involving the nAChR predicting the correct clockwise α-β-δ-α-γ arrangement of the five subunits. Models lacking this specific subunit arrangement were ignored and are not included in the discussion that follows.

**Figure 3.**
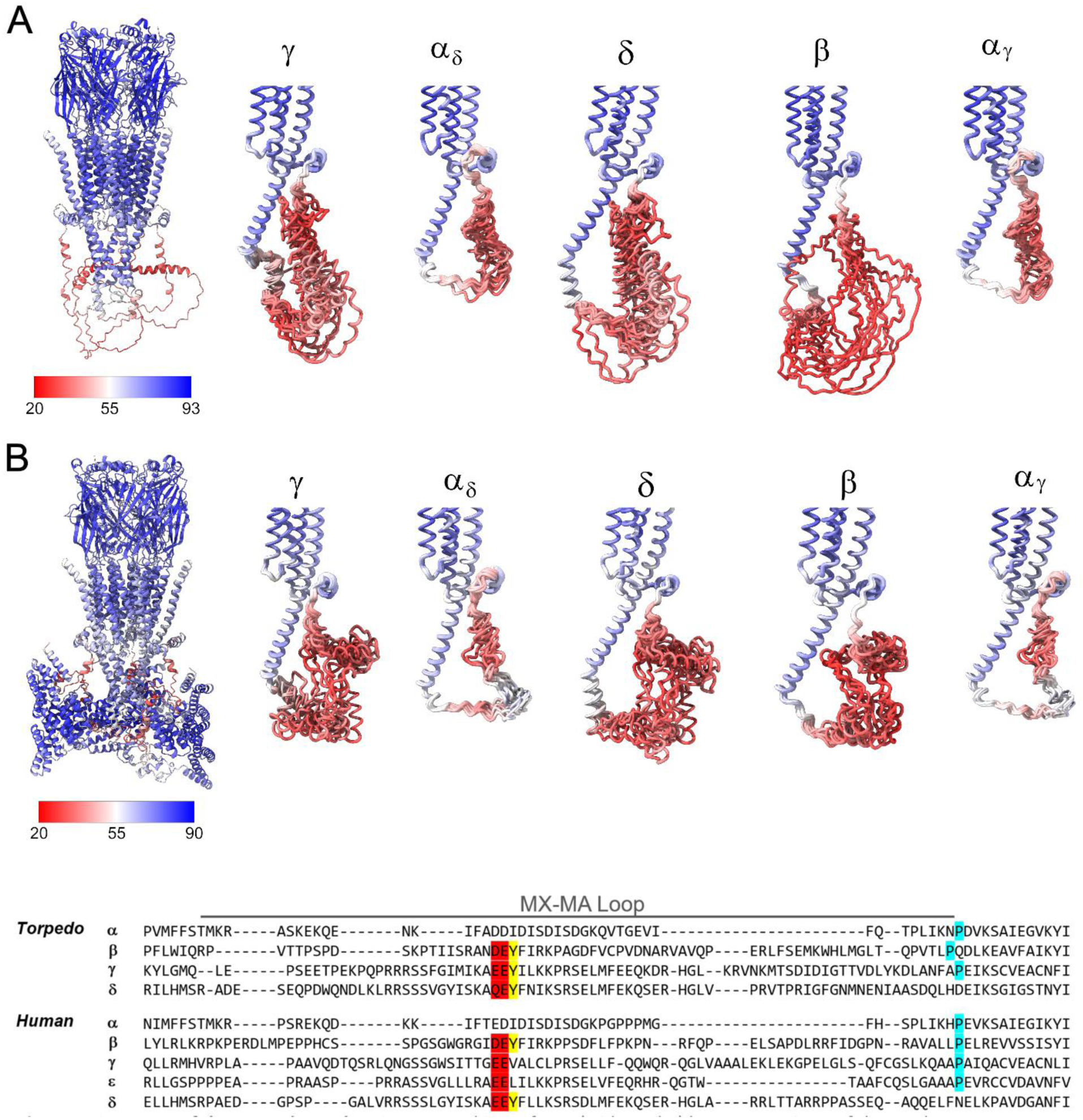
AlphaFold3 predictions of the Torpedo nAChR and rapsyn-bound complexes. A) Representative AlphaFold3 model of the Torpedo nAChR alone (left) and ensemble (15 models) of individual subunit predictions (right) colored by per residue pLDDT score from red (low confidence) to blue (high confidence). (B) Representative AlphaFold3 model of the 3:1, 3 PTR rapsyn–nAChR complex (left) and corresponding subunit ensembles (right) showing the predicted ICD conformations in the presence of bound rapsyn (15 models). Subunits are labeled γ, α_δ_, δ, β, and α_γ_, oriented identically across panels. Sequences of the MX-MA loop in each subunit of both the *Torpedo* and human nAChRs are shown below, with key phosphorylated tyrosine and acidic residues highlighted in yellow and red, respectively. Pro residues that define the start of each MA α-helix are highlighted in light blue.

**Figure 4.**
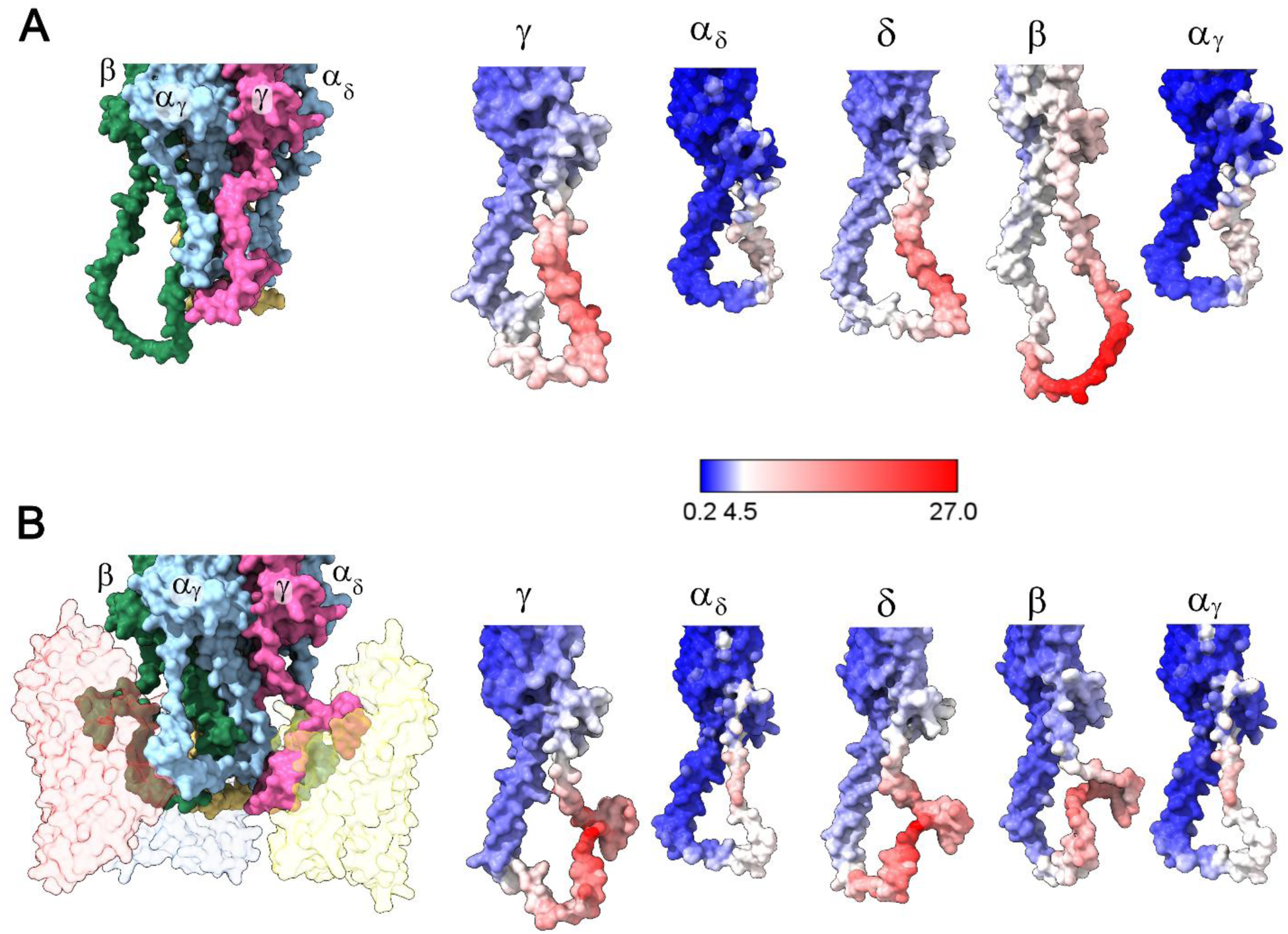
The structural variability of the ICD in ensemble predictions of the nAChR and the rapsyn–nAChR complex. **(A)** A surface side view of the average structure from ten random AlphaFold3-predicted *Torpedo* nAChR models is presented on the left, with the averaged structure of each individual subunit ICD color-coded on the right to illustrate the per-residue standard deviation (Å) across the predicted ensemble of structures from blue (0.2 Å, low variability) to red (27 Å, high variability). The individual subunits (γ, α_δ_, δ, β, and α_γ_) are each shown in a similar orientation to illustrate predicted differences in ICD flexibility across the receptor. **(B)** A surface side view of the average structure from ten AlphaFold3-predicted models of the nAChR with three bound rapsyns is shown on the left, with the averaged structure of each subunit ICD color-coded on the right to illustrate by per-residue standard deviation (Å) across the predicted ensemble of structures. On the left, rapsyn is shown as transparent surfaces with the transparent colors matching those in Fig. 6. Rapsyn is hidden in the individual subunit views on the right to highlight the conformational path of each polypeptide backbone in the predicted complex. All averages and standard deviations were computed after per-chain rigid-body alignment, in which each subunit and/or rapsyn was independently aligned using the N, C_α_, and C backbone atoms. The resulting SD values therefore represent internal conformational variability within each subunit across the predicted conformational ensemble rather than global rearrangements within the pentameric nAChR. Note that the ensemble of predictions for each subunit is shown in Fig. 3.

For each structure or complex, we typically generated between 15 and 25 models, initially evaluating each prediction using the predictive template modeling (pTM) score, the interface predictive template modelling (ipTM) score, the predicted Local Distance Difference Test (pLDDT), and the Predicted Aligned Error (PAE) matrix (Figs. 3 & S7-S8). The pTM score reflects the probable accuracy of the global structure, with pTM values above 0.5 suggestive of a high probability correct global fold. pTM values below 0.5 indicate low confidence in the predicted models. The ipTM values reflect the accuracy of the predicted interfaces between subunits within the model, with values above 0.8 corresponding to a high confidence in the predicted interfaces and values in the intermediate 0.6–0.8 range indicating only moderate confidence with potential structural inaccuracies (Evans et al., 2022; Jumper et al., 2021; Xu & Zhang, 2010; Zhang & Skolnick, 2004). The per-residue confidence metric, pLDDT, is used to assess local backbone confidence, with values above 70 indicating well-defined regions (Guo et al., 2022; Mariani et al., 2013). The PAE matrix provides complementary information on the relative positioning of domains and subunits, ranging from roughly 0 to 30 Å, with lower PAE values corresponding to smaller error ranges and thus higher confidence (Guo et al., 2022).

While these metrics provide a sense of the *probable accuracy* of the generated models, there are no absolute boundaries between “correct” and “incorrect” predictions (Edmunds et al., 2024). For example, pTM values higher than 0.5 provide a relative, not an absolute, measure of *potential* structural accuracy. In this context, rather than overinterpreting specific details of each structure or complex, we searched for recurrent architectural features that could be independently verified using available structural and biochemical data. We focus here on features that *visualize* verifiable constraints regarding the interactions between rapsyn and the nAChR.

### Models of the nAChR

Across the 15 models with the correct subunit arrangement, AlphaFold3 predicts with good confidence the overall structure of the *Torpedo* nAChR (pTM = 0.70-0.72) and with intermediate confidence the interfacial contacts (ipTM = 0.69-0.71) (Figs. 3, 4). These predicted structures superimpose well onto our ACh-bound cryo-EM structures and are reflective of the di-liganded state. On a per-residue basis, the ECD, TMD, and a major portion of the MA α-helices are predicted with pLDDT scores (70 to >90) ranging from high to very high confidence. The C-termini of the two α subunit M4 α-helices and both the immediate post-MX and pre-MA regions in all five subunits are predicted with pLDDT scores in the lower confidence 50 to 70 range. In contrast, the intervening MX-MA loops were consistently modeled with pLDDT scores of less than 50. These confidence levels were mirrored in the PAE matrices, which show inter-domain alignment errors of 1-4 Å for the ECD, TMD, and MA α-helices. The alignment errors increase to 4–6 Å for the α subunit M4 C-termini and both the immediate post-MX and pre-MA segments of each subunit (Fig. S8). The alignment errors increase further to 6–11 Å for the loops connecting these immediate post-MX and pre-MA structures. The combination of low prediction confidence and multiple suggested conformations for most of the MX-MA loops (Figs 3 & S7-S8), along with the absence of cryo-EM density in these regions in all *Torpedo* nAChR structures solved to date, support the idea each MX–MA linker naturally exhibits a flexible tertiary structure in the absence of stabilizing binding partners, such as rapsyn. To illustrate the predicted conformation of each ICD, we present both the ensemble of predictions colored by per residue pLDDT score (Fig. 3A) and an average of each ensemble colored by the per-residue standard deviation (Å) across the predicted ensemble of structures (Fig. 4).

The AlphaFold3 predictions are complementary to our refined structure of the ICD in that they reproduce experimentally observed features while extending the structures across the intrinsically disordered stretches of the MX-MA loops to form continuous ICD trajectories. In fact, the AlphaFold3 predictions accurately reproduce many subtle experimentally observed features in the new ICD structure, even though these features were not predicted with high confidence (Fig. 1B, C). Specifically, the predictions capture (1) the sharp and extended downward turn of the polypeptide backbone following MX; (2) pre-MA segments that follow a trajectory roughly parallel to the membrane surface in all subunits except δ; (3) the relative positioning of the pre-MA sequences at the apex of the MA inverted cone; (4) an abrupt bend in the pre-MA region of the α, β, and γ subunits as each polypeptide joins MA; (5) a shallower trajectory in the δ subunit as it joins MA; and (6) the N-terminus of each MA α-helix defined by a conserved proline residue in α, β, and γ. The strong correlations between observed structural features and the AlphaFold3 predictions lend confidence to the insight derived from the predicted structures. Importantly, the AlphaFold3 predictions, together with the constraints imposed by our new cryo-EM structures, permit us to visualize several *unequivocal* features that ultimately frame how the ICD loops of the nAChR interact with rapsyn (Figs. 3, 4).

First, both the AlphaFold3 predictions and our extended post-MX and pre-MA structures position the disordered ICD loops at the sides of the inverted cone formed by the five MA α-helices where they sterically shield MA from direct interactions with rapsyn and/or other cytoskeletal molecules (Figs 2 & 3). The lateral ICD positioning of the rapsyn binding sites is also supported by cryo-ET maps of *Torpedo* post-synaptic membranes (Figs 6 & S9).

Second, the ensemble of predictions shows that although the MX-MA loops from the β, γ, and δ subunits are each sufficient to form a flexible aqueous-exposed surface, each is not long enough to form an independent globular domain (Figs. 3, 4). Among these three ICDs, the β subunit is predicted with the greatest conformational variability. This variability may be functionally important, as it aligns with the unique predicted role of the β subunit in binding to both the principal face of one rapsyn molecule and the complementary face of another in the 3:1 nAChR-rapsyn complex. In fact, the 3:1 rapsyn-nAChR binding stoichiometry implied by cryo-ET maps of *Torpedo* postsynaptic membranes requires such a dual binding interface for one subunit ICD (see Fig. 6 and below). These predictions visualize that the MX-MA loops of the β, γ, and δ subunits act as flexible but spatially constrained scaffolding elements for adaptive binding to rapsyn and/or other cytoskeletal proteins.

Third, although the α subunit ICD is long enough to connect MX to MA, its shorter length leads to less conformational adaptability, than found in the longer MX-MA loops from the β, γ, and δ subunits, and thus less capacity to adapt its structure to fold around a bound rapsyn. In fact, the α subunit is predicted to extend down from MX prior to turning abruptly to join up with the roughly horizontal pre-MA sequence that is both predicted in the AlphFold3 models and defined in the ICD structure, as noted above. The result is a roughly V-shaped α-helical fold connecting MX to MA across all predicted models (Figs. 3, 4). Incredibly, this V-shape fold is predicted to bind to a complementary V-shaped groove on rapsyn in rapsyn-nAChR complexes (Fig. 6; see below). The predictions simply visualize that the shorter α subunit MX-MA loop must act as relatively static binding scaffold element with limited capacity to adapt its structure for rapsyn binding.

Fourth, the longer ICD loops of the β, γ, and δ subunits invariably occupy positions that sterically restrict access to the α subunit ICD loop, further supporting the contention that the β, γ, and δ form the primary interfaces for rapsyn binding. This arrangement aligns with the positions of three key phosphorylated tyrosine residues implicated in agrin-mediated *Torpedo* nAChR clustering (Huganir et al., 1984; K. Wagner et al., 1991; Borges & Ferns, 2001; Chrestia et al., 2023; Ferns, 2021).

### Rapsyn

In contrast to the nAChR, AlphaFold3 predicts the monomeric structure of the *Torpedo* rapsyn with high confidence (pTM = 0.92), with pLDDT values varying from 90–97 across the various TPR motifs and from and 67-69 across the more flexible RING-H2 domain. Notably, AlphaFold3 invariably predicts the same rapsyn tertiary fold with only subtle differences (Figs 5 & S7-S8), located mainly in the flexible C-terminus, in more than 150 predictions (both monomeric and oligomeric rapsyn, along with varying stoichiometric rapsyn-nAChR complexes). On the other hand, the software struggled to define a consistent mode of rapsyn self-association in oligomeric rapsyn structures, yielding multiple alternative dimer and trimer arrangements with substantially lower confidence (dimeric pTM = 0.50-0.56, iPTM = 0.14–0.18; trimeric pTM = 0.38–0.40, iPTM = 0.15–0.17). The diversity of low-confidence dimer and trimer poses may reflect the intrinsic binding promiscuity of rapsyn and its broad binding potential toward multiple protein partners, consistent with its established role as a postsynaptic scaffold (Apel & Merlie, 1995). Regardless, given these low prediction scores and the variability of the predicted interaction interfaces, we did not analyze further the potential oligomeric structures.

**Figure 5.**
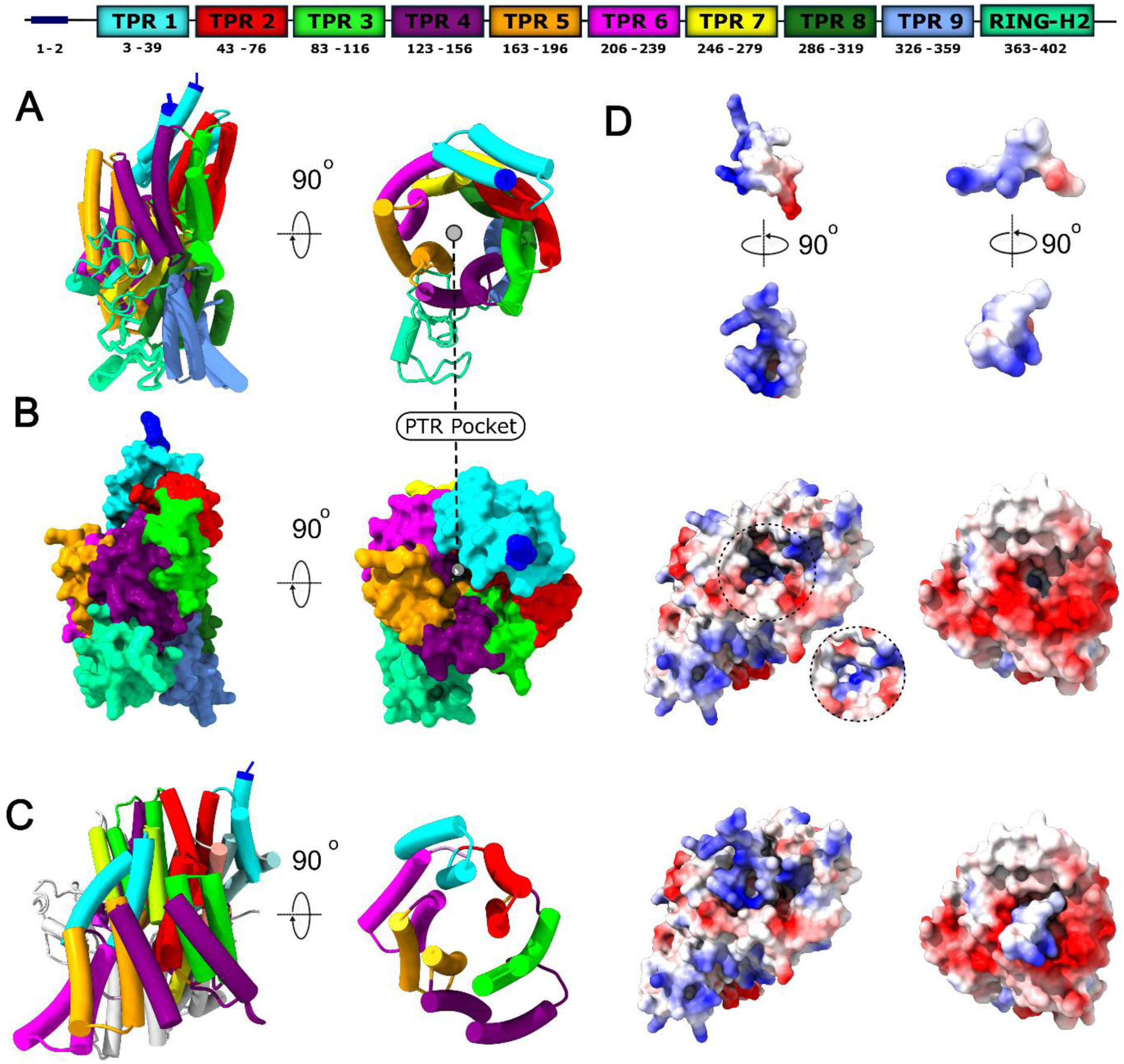
Comparison of the predicted model of rapsyn and portion of its bound MX-MA loop peptide ligand to that of the homolog, Pex5 (PDB: 9FH0). Side (left) and top-down (right) views of rapsyn are shown in both cartoon (α-helices as tubes) **(A)** and surface **(B)** representations with each TPR motif color-coded to match the sequence schematic in the top row. (**C)** Similar side and top-down views of the homolog, Pex5, shown in a cartoon representation, with the six aligning TPR motifs in Pex5 color-coded as in (A). **(D)** Surface representations of the electrostatic potentials of the predicted models of rapsyn and rapsyn-bound to a small, phosphorylated portion of the MX-MA loop from the β subunit (left column), along with similar views of Pex5 and Pex5 bound to a portion of its peptide ligand (right column), with blue and red indicating regions of positive and negative potential. The top row shows two views of an isolated portion of each bound peptide, with the lower of these two views corresponding to that of the same region of the bound peptide in the bottom row of (D). Views of rapsyn/Pex5 both without (middle row) and with a portion of its bound peptide are shown in the middle and bottom rows, respectively. The positive electrostatic surface potential deep in the pocket is shown with direct lighting, for clarity, in the inset.

In each of the rapsyn models, the predicted structure contains nine, instead of the originally proposed seven, TPR motifs, along with a globular RING-H2 domain (Frail et al., 1988). The eighth and ninth TPR motifs replace what had previously been predicted to be a coiled-coil domain (Ramarao & Cohen, 1998; see, however, Ponting & Phillips, 1996). Nine TPR motifs are also predicted using the bioinformatics tools TPRpred and HHpred (Gabler et al., 2020; Karpenahalli et al., 2007; Zimmermann et al., 2018) (Fig. 5). The nine TPR motifs form ∼1.5 turns of a right-handed super-helix. The full turn of the superhelix creates a concave surface with a shallow electropositive reservoir at its base, referred to here as the phospho-tyrosine reservoir, and a deep narrow pocket, referred to here as the phospho-tyrosine pocket, that extends below the phospho-tyrosine reservoir like a drain extending below a shallow sink. The RING-H2 domain projects away from the phospho-tyrosine reservoir and pocket of the super-helix. Electrostatic surface potentials show that the inner face of the superhelix is dominated by a strong positive electrostatic potential, partially resulting from three residues, Arg91, Lys95, and Arg164. These residues help create a charged environment compatible with phospho-tyrosine binding. Notably, the phospho-tyrosine pocket and surrounding electropositive region comprise numerous residues whose mutations lead to the CMS, underscoring the functional importance of this surface in nAChR–rapsyn interactions, as discussed below.

Although we have no structural constraints to assess the accuracy of the rapsyn prediction, the predicted super-helical arrangement of TPR motifs is consistent with known structures of other TPR-containing proteins (Bausewein et al., 2017; Chan et al., 2006; Gatto et al., 2000; Scheufler et al., 2000). In the AlphaFold3 rapsyn model, the TPR motifs form a tightly wound superhelix that exhibits a degree of curvature and packing similar to that observed in the peroxisomal receptor, Pex5, a protein with seven instead of nine TPR motifs (Stanley et al., 2007). The very high confidence predictions along with structural and electrostatic parallels between Pex5–peptide binding and the predicted rapsyn–phospho-tyrosine-nAChR binding reinforce the plausibility of the predicted rapsyn-nAChR complex model (Figs. 5C, 6, 7).

**Figure 6.**
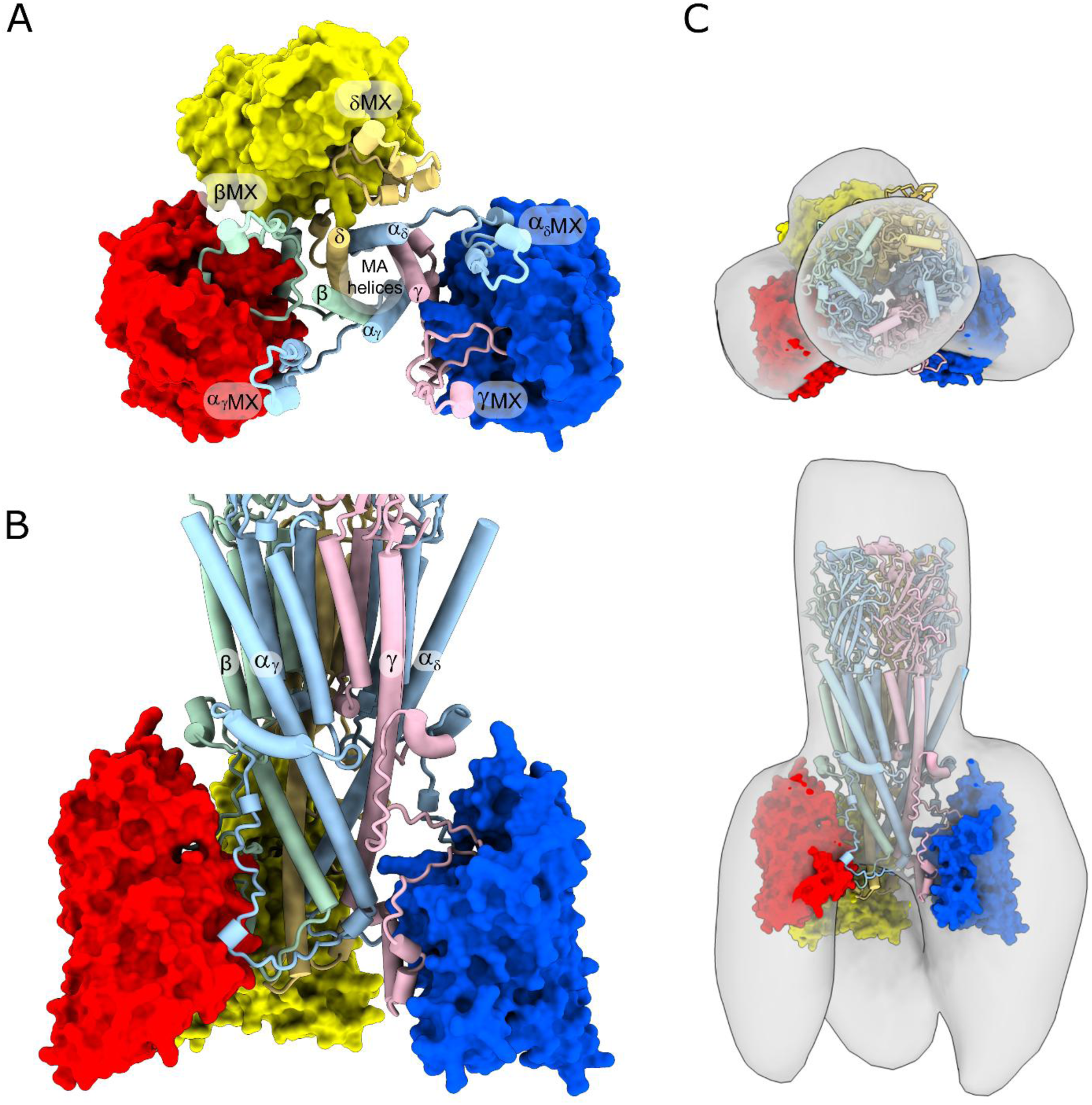
The predicted *Torpedo* rapsyn-nAChR complex and its fit into cryo-ET maps of *Torpedo* post-synaptic membranes. **(A)** A top-down view of a representative model of the rapsyn-nAChR complex showing the nAChR’s ICD, beginning in each subunit from a truncated MX α-helix and ending with a truncated MA α-helix, along with bound rapsyns presented as colored surfaces (red, yellow and blue). The color coding of the nAChR subunits is as in Fig. 1. **(B)** Side cartoon view of the same *Torpedo* rapsyn-nAChR complex focusing on the TMD and ICD of the nAChR. **(C)** Top down and side views of the predicted rapsyn-nAChR complex fit into cryo-ET density maps recorded from native *Torpedo* post-synaptic membranes (Zuber & Unwin, 2013).

**Figure 7.**
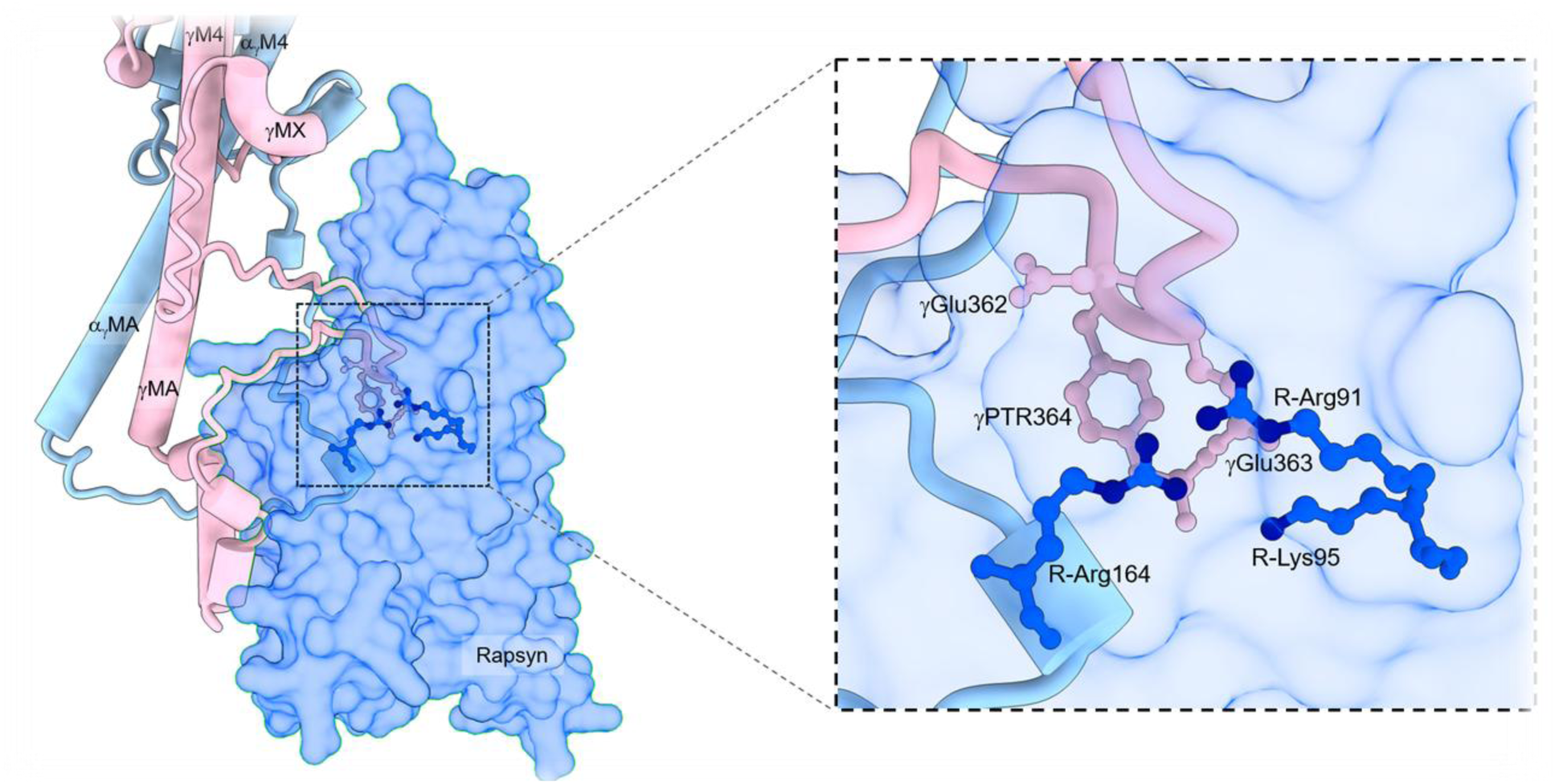
Predicted models of rapsyn sandwiched between the phosphorylated γ and the α subunits. Zoomed in side view of the rapsyn-nAChR complex showing a representative rapsyn (transparent blue surface) bound to the γ and α subunits (cartoon representation). The zoomed in view on the right highlights the predicted interactions between the phosphorylated tyrosine residue, γTyr364, and positively charged residues, Arg91, Lys95, and Arg164 in the PTR pocket of rapsyn.

### The rapsyn-nAChR complex

To explore how rapsyn interacts with the nAChR, we generated models of *Torpedo* rapsyn-nAChR complexes with stoichiometries ranging from one to more than three rapsyn molecules bound per nAChR pentamer, and with the nAChR in either a phosphorylated or an unphosphorylated state (between 10 and 25 models for each complex, >150 total models). We focused on the nAChR phosphorylated at the three sites, βTyr390, γTyr364, and δTyr393, known to facilitate interactions and thus clustering in *Torpedo* (Borges et al., 2008; Prömer et al., 2023). The predicted models of the rapsyn-nAChR complex have lower pTM/ipTM scores than the predicted models of the nAChR in the absence of rapsyn, with the scores progressively decreasing from an average of ∼0.64/∼0.61 for the 1:1 complex down to ∼0.52/∼0.49 for the 4:1 complex. This trend holds for both the phosphorylated and unphosphorylated states, although phosphorylation of the nAChR slightly increases the mean pTM/ipTM scores at each rapsyn stoichiometry. In all cases, local pLDDT values mirror those of the individual proteins: high confidence for the core regions of rapsyn and the nAChR (>70–90) and lower confidence for the ICD loops (<60). Interestingly, adding rapsyn (with or without nAChR phosphorylation) reduces local confidence in some regions of the nAChR that were predicted with high confidence in the absence of rapsyn, such as the M4 α-helices of all five nAChR subunits. For example, the pLDDT values for these five α-helices fall from 70-90 in the absence of rapsyn to below 70 in its presence. One possible explanation is that rapsyn binding alters the conformation of the M4 α-helices, potentially influencing their interactions with either the adjacent M1/M3 α-helices or lipids.

One remarkable feature of the predictions is that at stoichiometries of up to three rapsyn bound per nAChR, AlphaFold3 *always* predicts rapsyn binding with a pose that places its concave face, and thus both the PTR reservoir and PTR pocket, oriented towards the MX-MA loop of the β, γ or δ subunit. In this orientation, the N-terminal residue of rapsyn, and thus its myristylation site, is also *always* located close to the nAChR TMD and thus to the membrane surface. At low rapsyn-nAChR stoichiometries, the orientation of rapsyn relative to the long central nAChR axis varies slightly from one prediction to another with these variations in orientation correlating with variations in the predicted orientation of the binding partner MX-MA loop. As rapsyn numbers increase, the orientational variability decreases as binding sites on the ICD are filled and the MX-MA loops are forced into stricter binding poses. The predicted orientations of rapsyn are similar in both phosphorylated and unphosphorylated states, although phosphorylation alters the nature of how rapsyn interacts with each MX-MA loop (see below). Above the 3:1 rapsyn-nAChR stoichiometry, AlphaFold3 still predicts a complex with three rapsyns each bound in a similar orientation towards the β, γ, and δ subunit MX-MA loops, but with the additional rapsyn(s) bound in a range of poses, including poses where rapsyn binds to other rapsyn molecules or to non-ICD regions of the nAChR. Given that both the predicted rapsyn binding poses are similar for stoichiometries from one to three rapsyns per nAChR and the implied 3:1 stoichiometry of rapsyn binding at post-synaptic *Torpedo* membranes (Zuber & Unwin, 2013), we focus our discussion on the 3:1 rapsyn-nAChR complex.

In the 3:1 rapsyn-nAChR complexes, in both the phosphorylated and non-phosphorylated state, rapsyn is always (30 of 30 models) sandwiched between both a principal β, γ, or δ MX-MA loop and a complementary α_γ_, α_δ_ or β MX-MA loop, respectively. In most predictions (37 of 45 bound rapsyns) of rapsyn binding to the phosphorylated nAChR, the polypeptide backbone of β, γ, or δ extends down from MX to wrap into the PTR reservoir before spilling out of the reservoir to extend down the side of rapsyn near TPRs 4 and 5 (see Fig. 5A, 5B) for the predicted locations of the TPR motifs). Each β, γ, or δ MX-MA loop then wraps around the base of rapsyn prior to turning upwards to connect with each MA α-helix. In the remaining 8 of 45 bound rapsyns, the β, γ, and δ backbone adopts a similar overall pose, but instead of wrapping around the PTR reservoir, the chain extends directly down from MX along the side of rapsyn, again interacting with TPRs 4 and 5, before turning abruptly to wrap under rapsyn and rejoin MA. The resulting V-shaped pose is also observed in predictions (45 of 45) of rapsyn binding to the unphosphorylated nAChR. In all cases, the shorter MX-MA loops from α_γ_ and α_δ_ invariably fit into a V-shaped groove on the complementary face of rapsyn, which is formed by TPRs 6 and 7 (Figs. 5, 6 & 7). In addition to binding to the principal face of one rapsyn, the MX-MA loop from β interacts with the complementary face of a second rapsyn, although AlphFold3 is unable to predict the latter interactions with consistency.

One important feature of the predicted models is that both the complementary face of the δ-β rapsyn binding site and the principal face of the β-α_γ_ rapsyn binding site is formed by the same MX-MA loop from the β subunit. In fact, given the consistent binding of rapsyn between a longer more flexible β or γ subunit MX-MA loop and a shorter α subunit MX-MA loop, the relative positions of the subunits in the nAChR pentamer ultimately requires that the β subunit ICD forms a dual rapsyn binding interface in the 3:1 rapsyn-nAChR complex. The consequence is that with only the β subunit MX-MA loop separating the two bound rapsyns at the δ-β and β-α_γ_ sites, these two rapsyns are positioned closer to each other than to the rapsyn bound at the γ-α_δ_ site, thus leading to an asymmetric arrangement of rapsyn around the nAChR ICD. This predicted asymmetry fits well into the published low resolution cryo-ET volume maps obtained from *Torpedo* post-synaptic membranes (Zuber & Unwin, 2013) (Figs 6 & S9). The close fit between the predicted structures and the cryo-ET maps lends confidence to the hypothesis that up to three rapsyn bind to the nAChR ICD. The proposed sandwiching of each bound rapsyn between two subunit ICD loops provides a structural rationale for the asymmetry of rapsyn binding to the pentameric nAChR in post-synaptic *Torpedo* membranes, explains a wealth conflicting rapsyn-nAChR binding data regarding the subunit ICDs that interact with rapsyn (see Discussion) and provides a rationale as to why the ICDs of the β, γ, or δ subunits have evolved to be much longer than the ICDs of the α subunit.

Perhaps the most remarkable feature of the AlphaFold3 predictions is that the vast majority of rapsyns (37 of 45 bound rapsyns) are predicted to bind the phosphorylated nAChR such that the MX-MA loops of β, γ and δ extend into the phospho-tyrosine reservoir with each phosphorylated tyrosine, βTyr390, δTyr393, or γTyr364, and the preceding glutamate residue in each MX-MA loop projecting directly into the phospho-tyrosine pocket to interact with three positive residues, Arg91, Lys95 and Arg164 (Fig. 7). Moreover, many CMS-causing mutations are predicted either within or near the phospho-tyrosine binding pocket (Fig. 8) (see below). The remarkable convergence of prediction and biological data thus provides a plausible explanation of how phosphorylation of the nAChR enhances rapsyn-nAChR binding to promote nAChR clustering at the neuromuscular junction.

**Figure 8.**
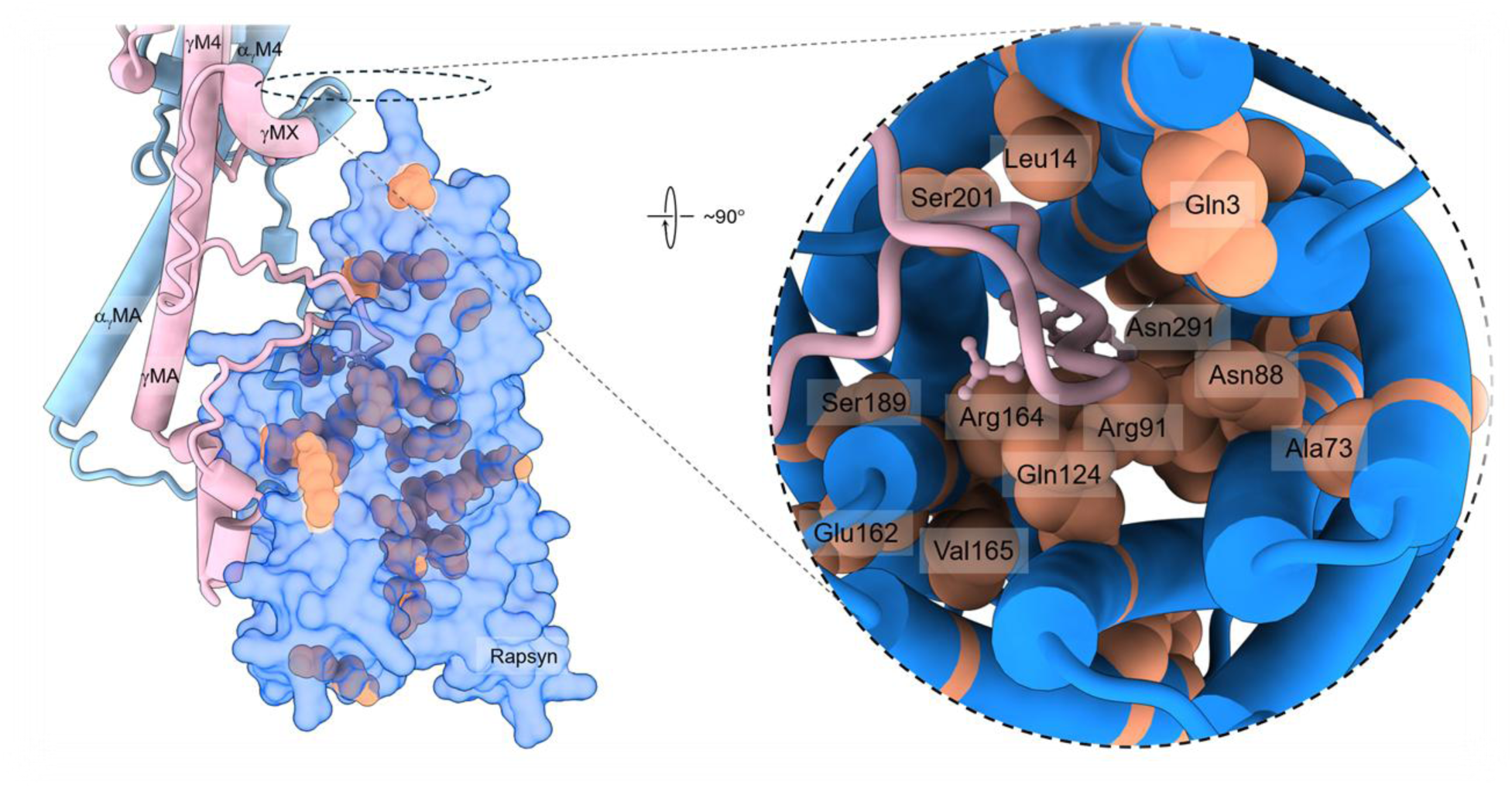
Predicted mapping of CMS-causing mutations in rapsyn. Side view the predicted model of rapsyn bound to the γ and α subunits with residues, whose point mutations lead to CMS, highlighted as light salmon-colored spheres. The zoomed in view on the right is a top-down view of the PTR reservoir and the PTR pocket.

### Mutations in rapsyn leading to CMS

As a further test of the predicted rapsyn-nAChR complex, CMS-causing mutations (Croxen et al., 1997; Engel et al., 2002; Liao et al., 2023a; Maselli et al., 2007; Milone et al., 2009; Müller, Baumeister, Rasic, et al., 2006; Müller, Baumeister, Schara, et al., 2006; Ohno et al., 2002; Xing et al., 2019) were mapped onto the predicted rapsyn structure bound to the nAChR (Fig. 8). Interestingly, the CMS-causing rapsyn mutations, R91L, R164C, and R164H, lie within the putative PTR binding pocket, where Arg91 and Arg164 are predicted to coordinate directly the negatively charged phospho-tyrosine residues in the β, γ and δ subunits of the *Torpedo* nAChR. Note that in the human nAChR, analogous phospho-tyrosines are located on the β and δ subunits and would be expected to interact with the human rapsyn in a similar manner. In contrast, the putative phosphorylated tyrosine in the γ/ε (fetal/adult) subunit is replaced with a hydrophobic Val/Leu residue, although the two preceding Glu residues could instead interact with the positively charged TPR reservoir and TPR pocket. Regardless, substitutions that neutralize or reverse the positive charges on Arg91 and/or Arg164 would be expected to reduce rapsyn’s affinity for both the phosphorylated β/δ and the unphosphorylated γ/ε subunits, thereby weakening complex formation. These predictions are consistent with the experimentally observed clustering deficits caused by these mutations (Cossins et al., 2006; Liao et al., 2023b).

The most prevalent CMS mutation, N88K, is predicted to be near the PTR pocket where it is located just above Arg91 (Figs 7 & 8). The change from a neutral to a positively charge lysine residue would introduce an additional positive charge in the PTR pocket, which could create electrostatic repulsion with Arg91, Lys95, and Arg164 to distort the PTR pocket and destabilize the phospho-tyrosine binding site. This interpretation aligns with the observation that N88K selectively disrupts rapsyn–nAChR association without impairing rapsyn self-association (Müller, Baumeister, Rasic, et al., 2006; Xing et al., 2019).

Several CMS-causing mutations are located close to or within the predicted RING-H2 domain. While this domain may not be strictly necessary for nAChR clustering (Bartoli et al., 2001), it still plays a role in the formation of the neuromuscular junction through its E3 ligase activity and its putative neddylation site, which may block ubiquitination to stabilize rapsyn (Bartoli et al., 2001; L. Li et al., 2016). The RING-H2 domain may also be an interaction partner for β-dystroglycan and utrophin, both important structural and functional proteins in the organization of the postsynaptic membrane (Bartoli et al., 2001).

Finally, other mutations are located throughout rapsyn at the interfaces between the α-helices that comprise individual TPR motifs or the interfaces between adjacent TPR motifs. Those in TPRs 1 and 2 are close to both the N-terminal myristylation site, which helps orient the complex towards the bilayer, and the PTR reservoir. Mutations in TPRs 7, 8, and 9 are on the face of rapsyn that projects away from the nAChR with the affected residues available for interactions with other proteins. Several of these residues are at the interface between the successive turns of the TPR super-helix. Mutations in these TPR motif interfacial residues may influence the folding of rapsyn to undermine its association with the lipid bilayer, the nAChR and/or other proteins.

Taken together, the predicted rapsyn-nAChR complex provides a structure-based rationale for how individual rapsyn mutations impair postsynaptic receptor clustering in CMS, thus lending support to the predicted rapsyn-nAChR model.

## Discussion

This work was motivated by the lack of defined structures of both rapsyn and the rapsyn-nAChR complex, which severely limits our understanding of agrin-mediated nAChR clustering at the neuromuscular junction. In this structural void, we used a focused mask to refine cryo-EM data leading to a map and thus a structure that has the most complete ICD density to date for the *Torpedo* muscle-type nAChR. Although two distinct conformations of the outermost transmembrane M4 α-helix were also resolved during refinement of the data, the main advance in this work is the improved ICD coverage, which places constraints on the conformational landscape adopted by each of the loops connecting the defined MX and MA α-helices. These constraints, along with published cryo-ET maps, then allowed us to validate a model of the nAChR-rapsyn complex that explains how agrin-induced phosphorylation of the nAChR promotes nAChR-rapsyn interactions to induce clustering at the neuromuscular junction. This model also that sheds light on how mutations in rapsyn impact rapsyn-nAChR interactions leading to CMS. The key findings of this work are as follows:

First, our new nAChR model reveals several previously undefined features of the ICD. The model shows that C terminal to each MX α-helix, the polypeptide chain in each subunit turns abruptly to follow a trajectory that extends roughly normal to the bilayer surface for up to 3 to 5 residues. No density is observed for a substantial stretch of each ICD immediately following this extension, consistent with other published structures of the muscle nAChR. Collectively, these structures confirm that this region of the MX-MA loop intrinsically lacks a well-defined tertiary structure in the absence of stabilizing binding partners, such as rapsyn. This intrinsically disordered stretch then follows a defined trajectory roughly parallel to the bilayer surface before turning abruptly upwards to join each MA α-helix. The short horizontal pre-MA sequences collectively cap the apex of the inverted cone-like structure formed collectively by the five MA α-helices. The abrupt turn to join MA in the α, β, and γ subunits is facilitated by a proline residue, with each proline defining the N terminus of each MA α-helix. The δ ICD lacks an analogous proline residue and undergoes a shallower turn to join up with MA. With a precise definition of the N terminal residue in each MA α-helix, we can now definitively place each phosphorylated tyrosine residue implicated in agrin-mediated *Torpedo* nAChR clustering, βTyr355, Tyr364 and δTyr372, in the intrinsically disordered stretch of the β, γ and δ subunits ICDs, respectively, implying that these regions of the polypeptide backbone are central to rapsyn-nAChR interactions (K. Wagner et al., 1991). The new structure also shows that the N and C boundaries, of each disordered MX-MA loop, project towards each other, thus placing them primarily laterally at the side of the ICD. As these structures serve as rapsyn binding sites, it can be concluded that rapsyn binds primarily adjacent to the inverted cone-like structure formed by the five MA α-helices, as opposed to near the apex of the inverted cone. The lateral positioning of the rapsyn binding sites is consistent with both electron microscopy/x-ray diffraction studies and cryo-ET maps of *Torpedo* post-synaptic membranes (Mitra et al., 1989; Zuber & Unwin, 2013).

Second, the predicted models of the nAChR reveal an ensemble of conformations that visualize the structural consequences of the different lengths of the MX-MA loops in the β, γ, and δ versus α subunits. The MX-MA loops in the β, γ, and δ subunits are long enough to adopt flexible conformations, but not long enough to form separate globular domains. The β, γ, and δ subunits MX-MA loops thus have the capacity to act as flexible but spatially constrained scaffolding elements for adaptive binding to rapsyn and/or other cytoskeletal proteins. The predictions also visualize that the extended lengths of the β, γ δ MX-MA loops are sufficient to restrict access to the MX-MA loops of the two α subunits, thus supporting the view that β, γ, and δ form the principal interfaces for rapsyn binding. In contrast, the two shorter α subunit MX–MA loops are constrained to being compact linkers with limited adaptability to bind rapsyn. In fact, AlphaFold3 reproducibly predicts that the MX-MA loops of the α subunits adopt a V shaped α-helical structure that is also predicted to bind a V-shaped groove on the surface of rapsyn.

Third, AlphaFold3 predicts with high confidence the monomeric structure of the *Torpedo* rapsyn. Although this model remains to be verified with experimental data, the predicted nine TPR motifs form a canonical right-handed super-helix consistent with known structures of other TPR-containing proteins, such as Pex5 (Banasik et al., 2024; Emmanouilidis et al., 2016), HSP organizing protein, Hop (Baindur-Hudson et al., 2015; Scheufler et al., 2000), and Tom70 (Chan et al., 2006; Fan et al., 2011), thus providing independent support that the modeled rapsyn fold represents a biologically realistic TPR framework. Notably, the predicted rapsyn super-helix generates a concave surface with a shallow electropositive reservoir, referred to as the phospho-tyrosine reservoir, at its base and a deep narrow pocket, referred to as the phospho-tyrosine pocket, that extends below the reservoir like a drain extending below a sink. Electrostatic surface potentials show that the inner face of the phospho-tyrosine pocket is dominated by a strong positive electrostatic potential, which is predicted to form a complementary charged environment for binding the phospho-tyrosine residue in the β, γ and δ subunits. Similarly, the homolog Pex5 binds a short, positively charged peptide within an electronegative reservoir and pocket that mimic the putative phospho-tyrosine reservoir and phospho-tyrosine pocket of rapsyn, respectively. The structural and electrostatic parallels between Pex5–peptide binding and the predicted rapsyn–phospho-tyrosine-nAChR binding reinforce the plausibility of the predicted rapsyn-nAChR complex model to provide a plausible explanation for agrin-mediated rapsyn binding to the nAChR.

Fourth, AlphaFold3 predicts a reproducible architecture for rapsyn binding to the nAChR that is consistent with considerable experimental data and that leads to a highly plausible model for agrin-mediated binding of rapsyn to the nAChR. All the predicted models have rapsyn binding with its phospho-tyrosine reservoir and phospho-tyrosine pocket facing the ICD in a manner that places its N-terminal myristylation site near the nAChR TMD and thus the lipid bilayer. This remarkably consistent feature is consistent with the important role the myristylation site plays in anchoring rapsyn to the membrane. Another key feature of the predicted rapsyn-nAChR complex is that although rapsyn binds primarily to the longer and more conformationally flexible β, γ, and δ MX-MA loops, which wrap around the bound rapsyn to form extensive contacts, rapsyn is always sandwiched between the MX-MA loops of two different subunits. Specifically, rapsyn is bound between β and α_γ_, γ and α_δ_, and δ and β. These predicted binding modes are consistent with experimental data showing that rapsyn can interact with every subunit ICD (Huebsch & Maimone, 2003; Maimone & Merlie, 1993), and validate prior studies suggesting that rapsyn binds at the interfaces between multiple subunits (Ferns, 2021). Moreover, the models explain experimental evidence that rapsyn interacts with the α subunit ICD even though the agrin-mediated phosphorylation sites driving rapsyn–nAChR complex formation are located exclusively on the β, γ, and δ subunits in *Torpedo* and on the β and δ subunits in mammalian nAChRs (Borges et al., 2008; Huebsch & Maimone, 2003; Huganir et al., 1984). The predictions suggest that while interactions with the phosphorylated β, γ, and/or δ subunits ultimately drive rapsyn–nAChR association, each α subunit ICD contributes a relatively static structural scaffold for complex formation.

A notable consequence of rapsyn binding between β and α_γ_, γ and α_δ_, and δ and β is that the two rapsyns located at the δ-β and β-α_γ_ interfaces are separated by the same β subunit MX-MA loop. This prediction highlights the dual role of the β subunit ICD as both a principal and a complementary rapsyn-binding face, which likely accounts for the repeated identification of the β subunit ICD as a primary site of rapsyn interaction (Borges et al., 2008; Gervásio et al., 2007; Huebsch & Maimone, 2003; Lee et al., 2009; Moransard et al., 2003). Notably, this binding motif also leads to asymmetry in rapsyn binding in that the rapsyns bound to the δ/β and β/α_γ_ interfaces are closer to each other than to the rapsyn bound to the γ and α_δ_ interface. The predicted 3:1 rapsyn-nAChR complex with asymmetric rapsyn binding fits well into published cryo-ET maps of *Torpedo* post-synaptic membranes (Zuber & Unwin, 2013), thus explaining the asymmetric arrangement of putative rapsyn molecules around the periphery of the nAChR in *Torpedo* post-synaptic membranes. The close fit between the predicted rapsyn-nAChR architecture and the experimental cryo-ET maps provides strong evidence in support of the predicted model. The binding of rapsyn to the MX-MA loops from adjacent subunits also echoes the multivalent binding interfaces seen in TPR-containing scaffolds such as Hop and PP5 (Baindur-Hudson et al., 2015, 2015; Das et al., 1998; Yang et al., 2005), where adjacent TPR grooves accommodate distinct peptide partners without major structural rearrangement. Thes experimental observations further reinforce the credibility of the rigid-body fits observed in the AlphaFold3 models.

Perhaps the most remarkable feature of the model is that the *Torpedo* rapsyn is predicted to adopt a fold that is both electrostatically and geometrically specialized for binding the phosphorylated ICD loops of the *Torpedo* β, γ, and δ subunits. The concave surface of the rapsyn superhelix, centered on Arg91, Lys95, and Arg164, is predicted to form a deep, positively charged phospho-tyrosine pocket that accommodates acidic and/or phosphorylated residues (Figs 5 & 7). In fact, the MX-MA loops of the β, γ, and δ subunits typically predicted to extend down into the phospho-tyrosine reservoir of rapsyn with their phosphorylated tyrosine residues projecting into the pocket to interact with Arg91, Lys95, and Arg164 (Figs 3 & 4). This arrangement follows the canonical TPR binding principle of concave-surface recognition via electrostatic complementarity, as exemplified by TPR proteins like Pex5 and Tom70, where amphipathic or acidic helices insert into polar grooves (Bausewein et al., 2017; Blatch & Lässle, 1999; Chan et al., 2006; Fan et al., 2011) (Fig. 5C, D). Rapsyn’s TPR superhelix, however, is unusually compact, forming a narrower and deeper pocket optimized for phosphate recognition. This structural logic extends to other TPR systems and parallels helix–loop–helix surface recognition modes seen in the TPR protein Fis1, which binds its partner Caf4 through an amphipathic helix wrapping into a shallow surface groove (Suzuki et al., 2003; Zeytuni & Zarivach, 2012; Zhang & Chan, 2007). In the AlphaFold3 rapsyn–nAChR models, the ICDs of the β, γ, and δ subunits similarly wrap an acidic helix–loop–helix element into the TPR groove, positioning the phosphorylated side chains toward the Arg/Lys triad. This hybrid mechanism - combining the binding pocket geometry of Tom70 with the helical clasp of Fis1 - offers a plausible structural basis for phosphorylation-dependent anchoring of rapsyn to the nAChR. Such a configuration would allow both high-affinity electrostatic binding and the flexibility required for dynamic receptor clustering at the postsynaptic membrane. Collectively, these analogues reinforce that rapsyn’s phosphate-selective pocket represents a specialized adaptation of a conserved TPR binding strategy.

These predictions also agree with biochemical evidence locating the three key phosphorylated tyrosine residues that mediate agrin-induced clustering of the *Torpedo* nAChR on the β, γ, and δ subunits (Huganir et al., 1984; K. Wagner et al., 1991; Borges & Ferns, 2001; Chrestia et al., 2023; Ferns, 2021). In *Torpedo* nAChR, these residues occur at βTyr355, γTyr364, and δTyr372 within the unstructured MX-MA loops (K. Wagner et al., 1991). The analogous β subunit tyrosine, βTyr390, in the mouse muscle nAChR undergoes agrin-induced phosphorylation to facilitate clustering (K. Wagner et al., 1991; Wallace et al., 1991) with the βY390F mutant in transgenic mice leading to smaller neuromuscular synapses, reduced numbers of AChR clusters, and fewer nAChRs within each cluster (Friese et al., 2007; Borges et al., 2008). Analogous phosphorylation sites are absent in the human in the γ/ε subunits, although in both cases binding to the PTR reservoir and PTR pocket of rapsyn could instead be facilitated by the preceding two glutamate/aspartate residues. Analogous phosphorylation sites are absent in both *Torpedo* and human α subunits, reinforcing the conclusion that the α subunit is unlikely to participate directly in agrin-mediated clustering (Ferns, 2021). The shorter α subunit ICD likely serves primarily as a binding facilitator rather than a high affinity rapsyn-binding site.

Note that although the presented model sheds light on the mode of interactions between rapsyn and the nAChR, several lines of evidence suggest that rapsyn self-association may mediate higher-order receptor clustering. Expression alone in *Xenopus* oocytes leads to rapsyn self-association at the plasma membrane, even in the absence of the nAChR (Froehner et al., 1990). AlphFold3 predicted with low confidence a wide range of different oligomeric rapsyn complexes from which we were unable to derive any reliable mechanistic insight. On the other hand, the ensemble of predicted interactions could suggest that rapsyn self association occurs through multiple modes of binding, a hypothesis consistent with the intrinsic binding promiscuity of rapsyn and its broad binding potential toward multiple protein partners, consistent with its established role as a postsynaptic scaffold (Apel & Merlie, 1995). Liquid-liquid phase separations may also facilitate the aggregation of rapsyn-nAChR complexes at post-synaptic membranes (Xing et al., 2021)

Fourth, the AlphaFold3 predictions provide a structural framework for interpreting the pathogenicity of rapsyn mutations implicated in CMS. Many CMS-causing mutations, including N88K, R91L, R164C, and R164H, lie within or adjacent to the putative phospho-tyrosine binding pocket, with Arg91 and Arg164 predicted to coordinate directly the negatively charged phosphor-tyrosines in the β and δ subunits of the human muscle nAChR. These residues may also coordinate the two preceding anionic residues in the fetal and adult γ/ε subunits. In addition, the most prevalent CMS mutation, N88K, is positioned just above Arg91 within TPR1. Substitutions that neutralize or reverse the local positive charge would be expected to reduce rapsyn affinity for phosphorylated nAChR subunits, thereby weakening complex formation. These predictions are consistent with the experimentally observed clustering deficits caused by these mutations (Cossins et al., 2006; Liao et al., 2023b). Furthermore, the introduction of a positive lysine residue at position 88 could create electrostatic repulsion with Arg91, Lys95, and Arg164 to distort the phospho-tyrosine binding pocket to destabilize phospho-tyrosine binding. This interpretation aligns with observations that N88K selectively disrupts rapsyn–nAChR association without impairing rapsyn self-association (Müller, Baumeister, Rasic, et al., 2006; Xing et al., 2019).

Multiple rapsyn missense mutations (e.g., N88K, R91L, R164H, S201N) also cluster within the TPR domains or at the interface between two turns of the rapsyn super-helix, particularly near the phospho-tyrosine binding pocket. These residues occupy regions essential for mediating electrostatic and hydrogen-bonding interactions with the receptor ICD. Consistent with this, structural modeling of the αβδ–rapsyn interfaces reveals that mutations within TPR4 may disrupt hydrogen bonding with acidic residues in the β subunit ICD, whereas N88K and R91L, both located in TPR1, likely destabilize the orientation of the N-terminal TPR lobe that interfaces with the δ subunit’s extended MA helix and adjacent loop. While these ICD elements are not directly resolved in the cryo-EM density, their positions are geometrically constrained by neighboring regions and align closely with the predicted rapsyn contact zones.

Finally, heterogeneity observed in our cryo-EM models for the α_δ_ M4 α-helix, manifested as distinct tight and tilted conformations, raises questions about both putative M4 dynamics and possible interference from the surrounding nanodisc. The α_δ_ M4 α-helix is uniquely affected, adopting two separable poses that coexist within the same particle ensemble. This asymmetry suggests either a subunit-specific functional role for M4 or a preferential interaction of the α_δ_ face with the nanodisc scaffold/lipid boundary. Given that the α_δ_ M4 lies at the periphery of the receptor, it is particularly susceptible to variations in lipid composition and curvature, which could modulate its orientation. It has likewise been suggested that some proteins prefer to adopt a conserved orientation within the MSP ring of a nanodisc, which could explain why α_δ_ is the subunit most consistently impacted (Dalal et al., 2024).

The presence of both tight and tilted M4 α-helices under desensitizing conditions suggests that these conformations correspond to distinct substates within the desensitized ensemble, possibly influenced by agonist occupancy. It was recently shown (Thompson et al., 2025) that agonist binding proceeds sequentially, with the α_γ_ site binding and capping first, priming the receptor for α_δ_ binding and activation. By analogy, in desensitizing conditions, α_δ_ may be the first site to unbind agonist, yielding a structural transition in which the close (tight) M4 could represent a pre-desensitized intermediate. On the other hand, both the tight- and tilted M4 models contain agonist densities in capped conformations, consistent with the possibility that the conformational heterogeneity reflects interactions with the membrane/nanodisc rather than complex gating transitions.

These observations highlight the potential influence of the nanodisc itself on local conformational equilibria, particularly for peripheral α-helices, such as M4, that interface directly with lipids. The shorter α subunit M4 α-helices may further increase susceptibility to nanodisc-induced tilting, although the consistent involvement of the α_δ_ M4 implies a conserved or geometrically favored pose within the nanodisc scaffold. Importantly, this effect emerges only at higher agonist concentrations (Rahman et al., 2022), suggesting that conformational changes during activation or desensitization may expose hydrophobic surfaces that interact more strongly with the surrounding lipid or nanodisc environment.

The presence of both a tight and tilted M4 conformation in the same sample challenges the notion that M4 tilting is a strict marker of gating state and instead may suggest a role for lipid or scaffold protein interactions (or interference) in M4 dynamics. M4’s location on the periphery of the protein makes it more susceptible to modulation by lipids (Baenziger & daCosta, 2013; Barrantes, 2004; Blanton & Cohen, 1994; Carswell et al., 2015; Hénault et al., 2015, 2015, 2019) and presumably, to interactions with the nanodisc (Dalal et al., 2024). The shorter α subunit M4 α-helices may further increase susceptibility to nanodisc-induced tilting, although the consistent involvement of the α_δ_ M4 implies a conserved or geometrically favored pose within the nanodisc scaffold. That this effect is only seen at higher agonist concentrations may indicate a combined effect wherein the M4 becomes more susceptible to nanodisc or lipid interference in the desensitized structure, perhaps as a result of conformational changes during activation.

## Conclusions

In summary, while AlphaFold has been widely applied to predict full-length protein structures or assign densities in large cryo-EM complexes, its use to refine previously unresolved domains within high-resolution experimental models remains uncommon. This study demonstrates the power of combining focused classification strategies with predictive modeling to clarify the structures of long-standing blind spots in cryo-EM models of ligand-gated ion channels. By integrating experimental and computational approaches, we generate a highly plausible model of the rapsyn-nAChR complex that is consistent with extensive structural and biochemical data, and that explains how rapsyn senses both sequence-specific docking motifs and post-translational cues to mediate selective and dynamic receptor clustering. Looking forward, future studies incorporating cryo-ET, integrative modeling, and lipid-specific nanodiscs and liposomes will be essential to dissect the interplay between receptor structure, scaffold proteins such as rapsyn, and the membrane environment. Such efforts will deepen our understanding of neuromuscular synapse formation, stability, and plasticity, and may ultimately inform strategies to target pLGIC dysfunction in neurodegenerative, psychiatric, and neuromuscular diseases.

## Materials and Methods

### nAChR purification and nanodisc reconstitution

The nAChR was purified from the electroplax tissue (Aquatic Research Consultants, CA) of the Pacific electric ray *Tetronarce californica* (NCBI: txid7787) using a bromoacetylcholine bromide-derivatized Affi-Gel 102 column (Bio-Rad) as previously described (Zarkadas et al., 2022), but with the following modifications. After the nAChR was eluted from the column, it was incubated with 15 mM DTT to break disulfide-linked pentamers, followed by purification on a Superose6 size-exclusion column, in the presence of 1.05mM soybean azolectin (Sigma).

Following size-exclusion, the monomeric form of the nAChR pentamer was incubated on ice, still in the presence of 1.05mM soybean asolectin with purified MSP2N2 at a 1:5 molar ratio, with sufficient total lipid to obtain a final 1:5:500 ratio of nAChR:MSP:Lipid. The sample was then dialysed five times (12-14 kDa MW cut-off) against 2L of tris dialysis buffer, with the buffers changed roughly every 12 h. Following dialysis, the MSP-reconstituted nAChR was further purified by size-exclusion chromatography, with the appropriate fractions pooled, concentrated to 0.65 mg/mL, snap frozen in liquid nitrogen and stored at −80°C.

### Cryo-EM Sample Preparation and Data Collection

The nAChR in asolectin-MSP2N2 nanodiscs was mixed with the megabody Mbc7HopQNbF3 (Uchański et al., 2021), which binds MSP2N2, not the nAChR, at a molar ratio 1:3 to overcome orientation bias of the nAChR on the cryo-EM grids. After incubating with 50µM ACh for 30 min on ice, 3.5µls of sample were deposited onto a glow-discharged (30 mA, 50s) Quantifoil Au/C R 1.2/1.3 grid, blotted for 6 s with force 0 at 8 °C and 100% humidity, and then plunge-frozen in liquid ethane using a Mark IV Vitrobot (Thermo Fisher Scientific).

Datasets were recorded on a Glacios electron microscope at the Institut de Biologie Structurale in Grenoble, France. Movies (40 frames) were recorded at a nominal magnification of 36,000 with SerialEM and a Gatan K2 Summit camera. The raw movies were imported to Cryosparc (Punjani et al., 2017), aligned, and summed. CTF estimation was calculated for the non-dose-weighted sums. Particles were autopicked with crYOLO (T. Wagner et al., 2019) using the general model for low-pass filtered images. Particle coordinates (∼2,200,000) were imported into Cryosparc, where all subsequent data processing was performed. Raw particle stacks were extracted using 256-pixel box sizes.

The dataset was subjected to multiple rounds of 2D classifications, *ab initio* reconstruction, and heterogeneous refinement (3 classes). The class showing the best nAChR ICD features (map integrity and isotropic viewing angle distributions) was selected and refined using non-uniform and local refinement (Punjani et al., 2020) to yield a consensus reconstruction. The reconstruction was refined to improve the view of the ICD using masks focused on that region. This strategy produced reconstructions at 3.04Å resolution from ∼139,000 particles, with well-defined ICD suggesting that additional residues could be modelled, however this map also presented ambiguous αδ M4 helix features, suggesting that two M4 poses were present within the same sample set.

The map was thus further processed for conformational heterogeneity at the α_δ_ M4 using 3D classification and 3D variability analysis (3DVA) (Punjani & Fleet, 2021) with focused masks encompassing the ICD and the M4-M1/M3 region of the αδ subunit. Subclassification of α_δ_ M4 density yielded two particle subsets: a compact α_δ_M4 conformation at 3.16 Å (∼48,000 particles) and a tilted α_δ_M4 conformation at 3.05 Å (∼90,000 particles). An additional 2.68 Å map was also generated and used to verify the position of the M4 residues in the model, but as it had a lower resolution of the ICD, it was not subjected to further processing.

### Model refinement and structure analysis

Preliminary models were docked into unsharpened maps in ISOLDE (Croll, 2018). Iterative manual rebuilding was carried out in Coot (Emsley et al., 2010) with real-space refinement in Phenix (Liebschner et al., 2019). Final models were validated against unsharpened maps using MolProbity (Williams et al., 2018).

### Alphafold3 structure generation

Full-length models of the *Torpedo* nAChR, *Torpedo* rapsyn, and rapsyn-nAChR complexes were generated using AlphaFold3. Predictions were assessed for the correct subunit assembly order, with models excluded based on incorrect assembly or inter-woven ICD loops. Correctly assembled structures were evaluated for quality using template modeling scores (pTM and ipTM). Regions of the extracellular domain (ECD), transmembrane domain (TMD), and MA helices were predicted with high confidence (ipTM ≥ 0.8), while the flexible regions between the MX and MA loops were predicted with lower confidence (ipTM 0.6–0.7). Predicted conformational rapsyn-nAChR ensembles were compared with the improved ICD cryo-EM maps to evaluate the plausibility of the modelled ICD–rapsyn interaction modes.

## Acknowledgments

We thank Eleftherios Zarkadas for the cryo-EM data acquisition and Hugues Nury for assistance with the ICD-focused refinement of the cryo-EM data and for feedback on the manuscript. We thank Guy Schoehn for establishing and managing the IBS-ISBG cryo-electron microscopy platform and for providing access, training and support. The cryo-EM data were recorded using the platforms of the Grenoble Instruct-ERIC center (ISBG; UAR 3518 CNRS-CEA-UGA-EMBL) within the Grenoble Partnership for Structural Biology (PSB), supported by FRISBI (ANR-10-INBS-0005-02) and GRAL, financed within the University Grenoble Alpes graduate school (Ecoles Universitaires de Recherche) CBH-EUR-GS (ANR-17-EURE-0003). The IBS-ISBG EM facility is supported by the Auvergne-Rhône-Alpes Region, the Fondation Recherche Medicale (FRM), the fonds FEDER and the GIS-Infrastructures en Biologie Sante et Agronomie (IBISA).

## Author contributions

JEB designed the research project. CH and JEB processed the cryo-EM data with CH performing the protein modeling. AlphaFold3 predictions were generated by MH and CH in consultation with JEB. CH, CJGT and JEB wrote the manuscript and prepared the figures.

## Data Availability

All atomic models and cryo-EM maps deposited in the Protein Data Bank and Electron Microscopy Data Bank will be released upon acceptance of the manuscript. AlphaFold3 predictions will be deposited in a public data bank.

