## Supplementary Data File for "An extended structure of the intracellular domain of the Torpedo nicotinic acetylcholine receptor and its proposed interactions with rapsyn"

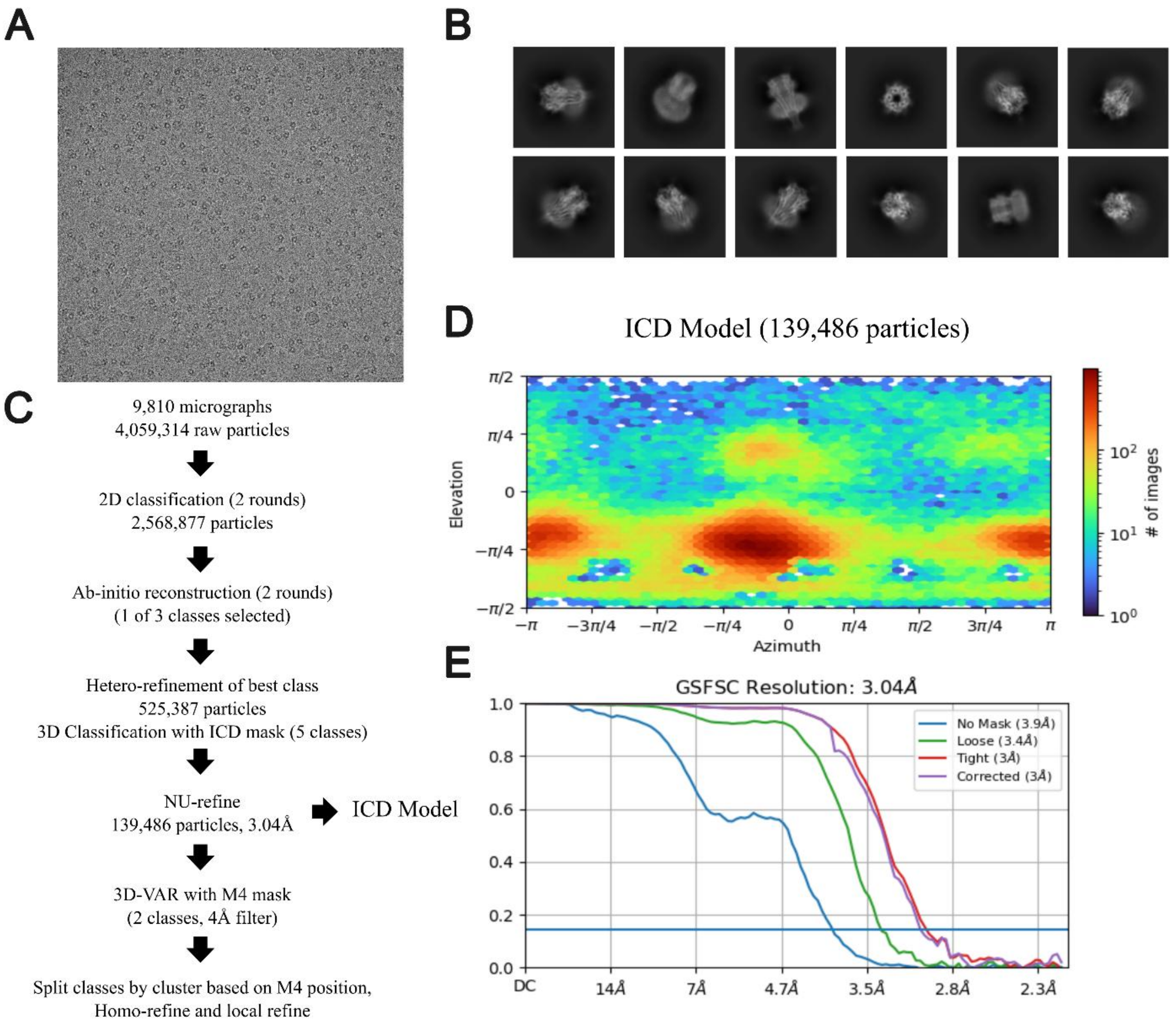


**Figure S1 Cryo-EM data processing and reconstruction of the ICD-focused *Torpedo* nAChR map.** (A) Motion- and CTF-corrected cryo-EM micrograph from the dataset collected at 50 μM ACh. (B) Representative 2D class averages of the *Torpedo* nAChR. (C) Image-processing workflow showing particle selection, two rounds of 2D classification, ab-initio reconstruction, heterogeneous refinement, and 3D classification using an ICD mask. Non-uniform refinement of the selected class (139,486 particles) produced the ICD-focused map at 3.04 Å resolution. Subsequent 3D variability analysis (3D-VAR) with an M4 mask separated the data into two classes corresponding to Tight and Tilt M4 conformations. (D) Angular distribution of particle orientations for the ICD model. (E) Gold-standard Fourier shell correlation (GSFSC) curves for the unmasked, masked, and corrected maps showing a final global resolution of 3.04Å. Additional FSC curves (no mask, loose mask, tight mask, and corrected masked) are shown to illustrate the effect of masking on resolution estimation. Displaying all curves highlights how focused masking of the ICD region progressively improves map correlation and supports the reported resolution values.

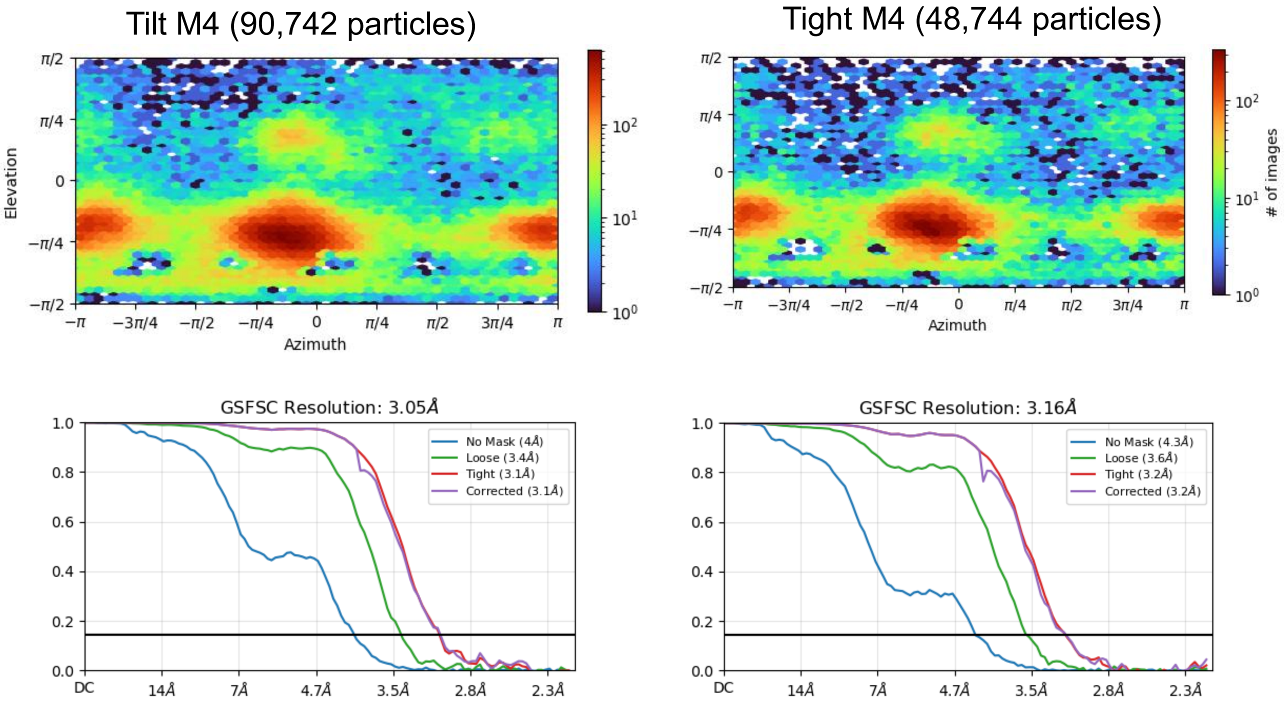


**Figure S2 Cryo-EM data processing for the $\alpha_{\delta}$M4-Tight and Tilt and models.** Angular distribution plots and corresponding gold-standard Fourier shell correlation (GSFSC) curves for the M4-tilt (left) and M4-tight (right) reconstructions. Each map displays isotropic angular sampling (top panels) and overall resolutions of 3.05 Å and 3.16 Å (bottom panels), respectively. Additional FSC curves (no mask, loose mask, tight mask, and corrected masked) are shown to illustrate the effect of masking on resolution estimation. Displaying all curves highlights how focused masking of the ICD region progressively improves map–map correlation and supports the reported resolution values.

A

B

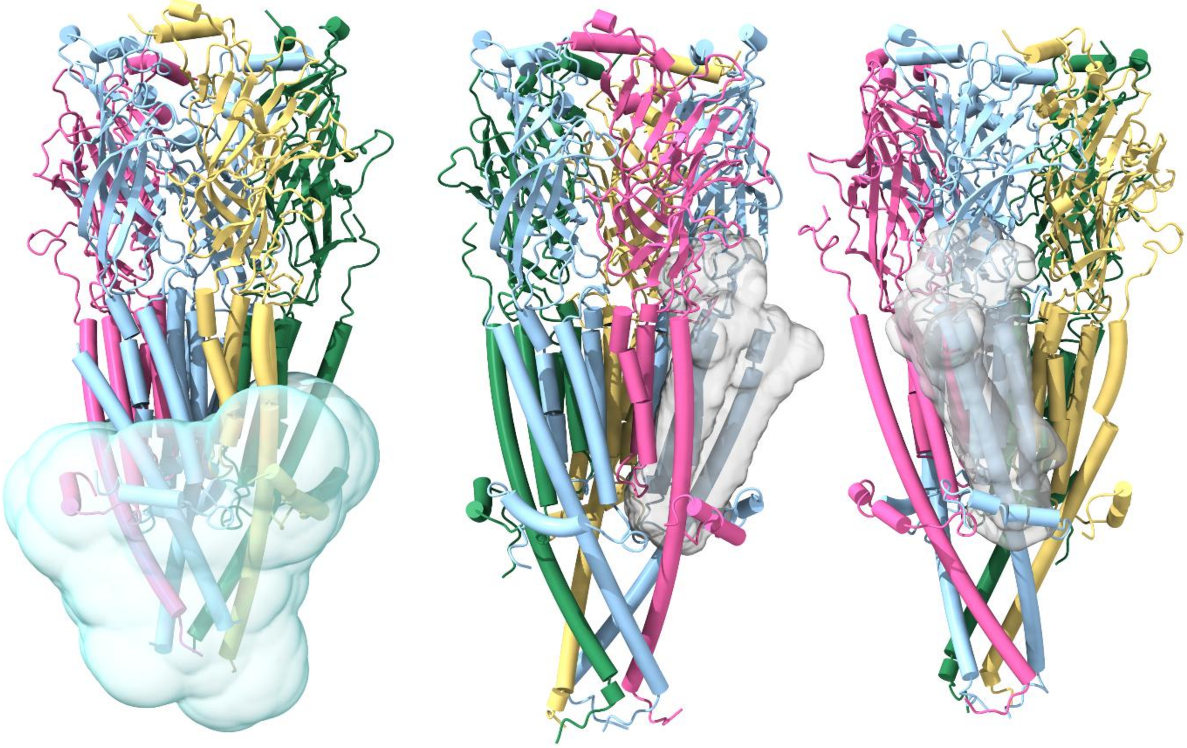

**Figure S3. Focused masks** used in A) ICD 3D classification and Hetero-refinement, and B) 3D-Variability analysis of the $\alpha_\delta$M4-M1/M3.

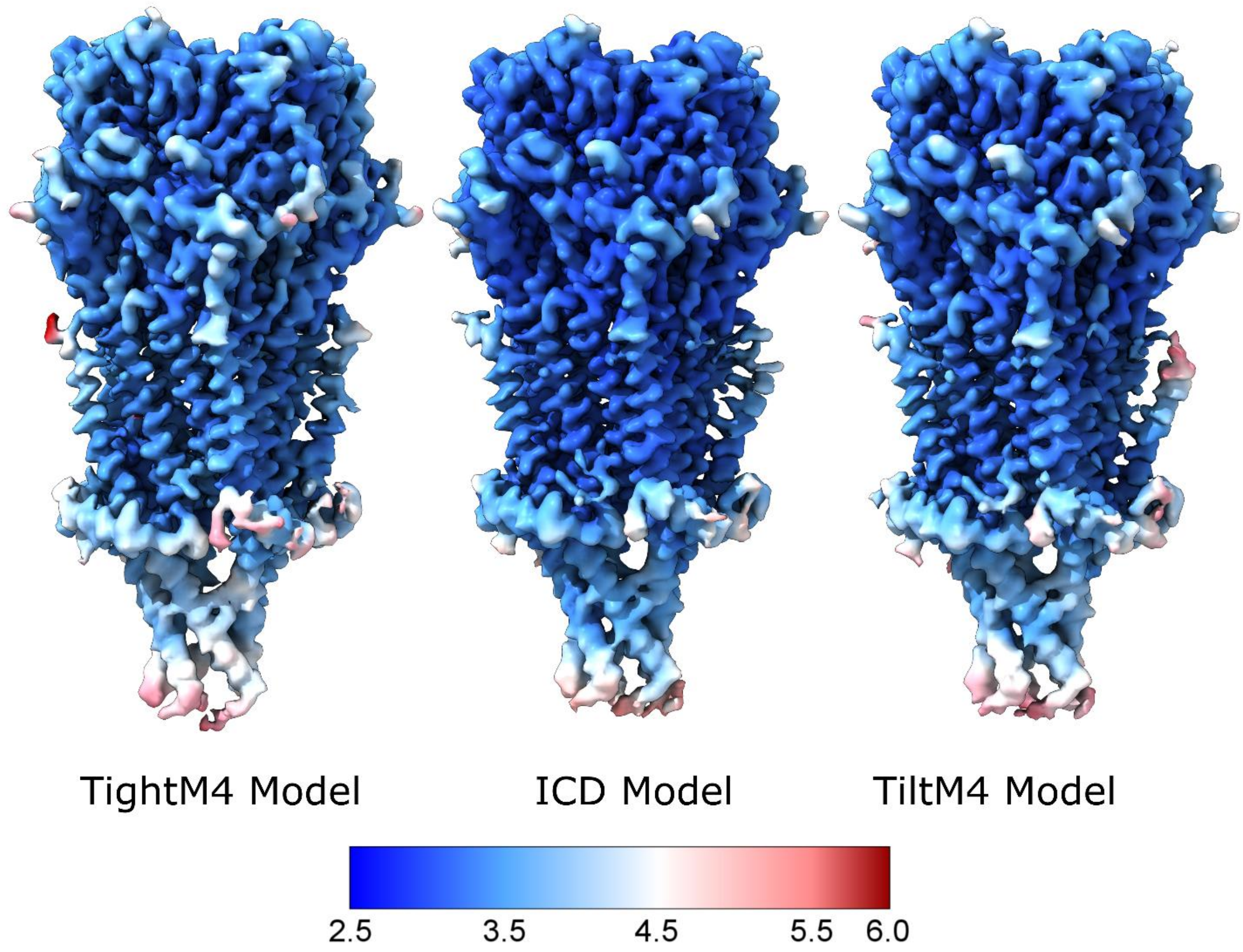


**Figure S4 Cryo-EM maps of the *Torpedo* nAChR showing the ICD-focused reconstruction and M4 conformational heterogeneity.** Cryo-EM density maps of the TightM4, ICD, and TiltM4 reconstructions are shown, colored by local resolution as indicated by the scale bar (2.5–6.0 Å). The maps correspond to models refined using focused classifications on the ICD and M4 regions.

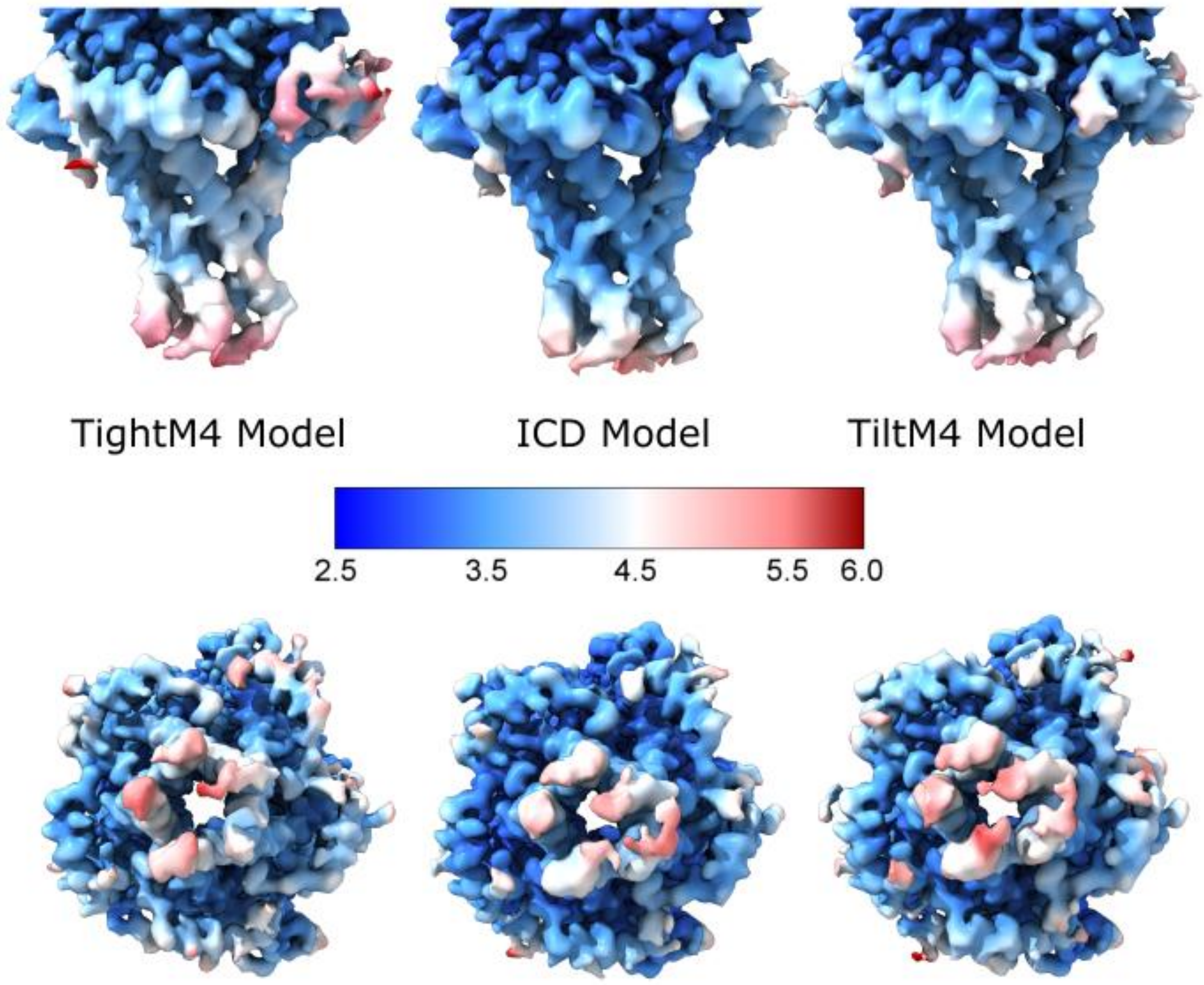


**Figure S5 Views of the intracellular domain (ICD)** from the $\alpha_{\delta}$M4-Tight, ICD, and $\alpha_{\delta}$M4-Tilt reconstructions, colored by local resolution (2.5–6.0 Å). The maps correspond to models refined using focused classifications on the ICD and M4 regions.

```
Alpha-Gamma
              MX    Post-MX                 ICD Loop                              MA
             321                                                                   380
Torpedo     PNVMFFSTMKRASKEKQENKIFADDIDISDISGKQVTGEVIFQTPLIKNPDVKSAIEGVK
ICD Model   PNVMFFSTMKRASKE-----------------------------TPLIKNPDVKSAIEGVK
9e3e        PNVMFFSTMKRP-------------------------------------DVKSAIEGVK
7ql5        PNVMFFSTMKR-----------------------------------------SAIEGVK

Alpha-Delta
              MX    Post-MX                 ICD Loop                              MA
             321                                                                   380
Torpedo     PNVMFFSTMKRASKEKQENKIFADDIDISDISGKQVTGEVIFQTPLIKNPDVKSAIEGVK
ICD Model   PNVMFFSTMKR--------------------------------QTPLIKNPDVKSAIEGVK
9e3e        PNV-----------------------------------------------------GVK
7ql5        PNV-----------------------------------------------------GVK

Beta
              MX     Post-MX                              ICD Loop                                              MA
             327                                                                                                  410
Torpedo     PPFLWIQRPVTTPSPDSKPTIISRANDEYFIRKPAGDFVCPVDNARVAVQPERLFSEMKWHLNGLTQPVTLPQDLKEAVEAIKY
ICD Model   PPFLWIQRPVTTP----------------------------------------------------------VTLPQDLKEAVEAIKY
9E3E        PPFLWIQRP-----------------------------------------------------------------QDLKEAVEAIKY
7QL5        PPFLWIQR-----------------------------------------------------------------------VEAIKY

Gamma
              MX   Post-MX                                   ICD Loop                                                   MA
             330                                                                                                              427
Torpedo     PKYLGMQLEPSEETPEKPQPRRRSSFGIMIKAEEYILKKPRSELMFEEQKDRHGLKRVNKMTSDIDIGTTVDLYKDLANFAPEIKSCVEACNF
ICD Model   PKYLGMQ----------------------------------------------------------------DLANFAPEIKSCVEACNF
9E3E        PKYLGM-----------------------------------------------------------------------APEIKSCVEACNF
7QL5        PKYLGM--------------------------------------------------------------------------------EACNF

Delta
              MX   Post-MX                                   ICD Loop                                                MA
             334                                                                                                         424
Torpedo     LPRILHMSRADESEQPDWQNDLKLRRSSSVGYISKAQEYFNIKSRSELMFEKQSERHGLVPRVTPRIGFGNNNENIAASDQLHDEIKSGID
ICD Model   LPRILHMSRAD-------------------------------------------------------------------- DQLHDEIKSGID
9E3E        LPRILHMSE-------------------------------------------------------------------------DEIKSGID
7QL5        LPRILH------------------------------------------------------------------------------ --IKSGID
```

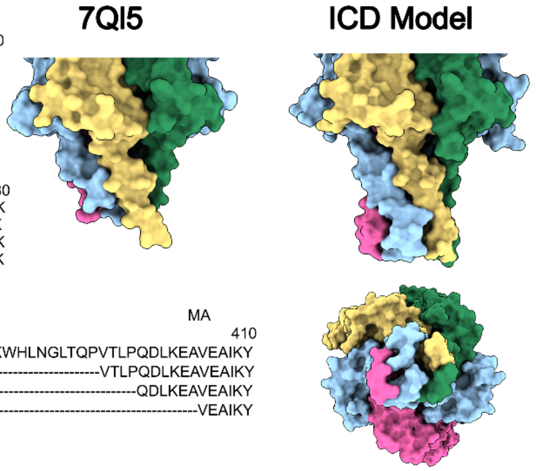


**Figure S6 Comparison of modeled ICD regions among *Torpedo* nAChR subunits.** Sequence alignments of the MX–MA regions from each Torpedo nAChR subunit are shown, comparing the full *Torpedo* sequence with the ICD-focused model, ACh-bound structure (PDB: 9E3E), and nicotine-bound structure (PDB: 7QL5). Dashes indicate residues absent from the corresponding model. Right, surface representations of the 7QL5 (left) and ICD (right) models showing the additional density corresponding to the extended ICD regions. Subunits are colored as in Fig. 1.

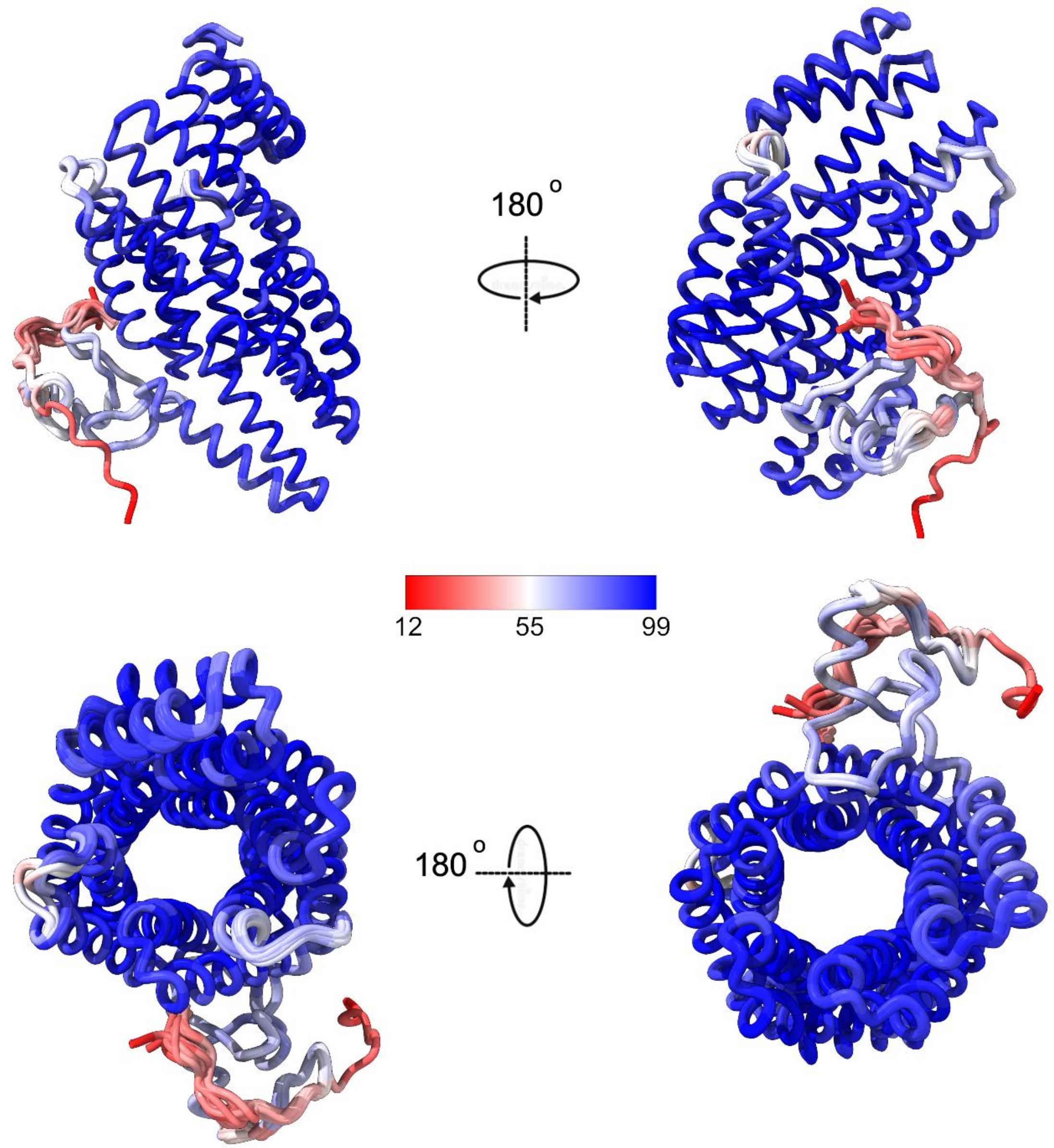


**Figure S7 Ensemble of AlphaFold3-predicted rapsyn structures.** Overlay of 15 predicted rapsyn monomer models from AlphaFold3, shown in two opposing side views (top) and corresponding top-down views (bottom). Models are colored by per-residue pLDDT confidence score from red (low confidence) to blue (high confidence). All models adopt a consistent overall fold with well-defined tetratricopeptide repeat (TPR) architecture and variable N- and C-terminal regions.

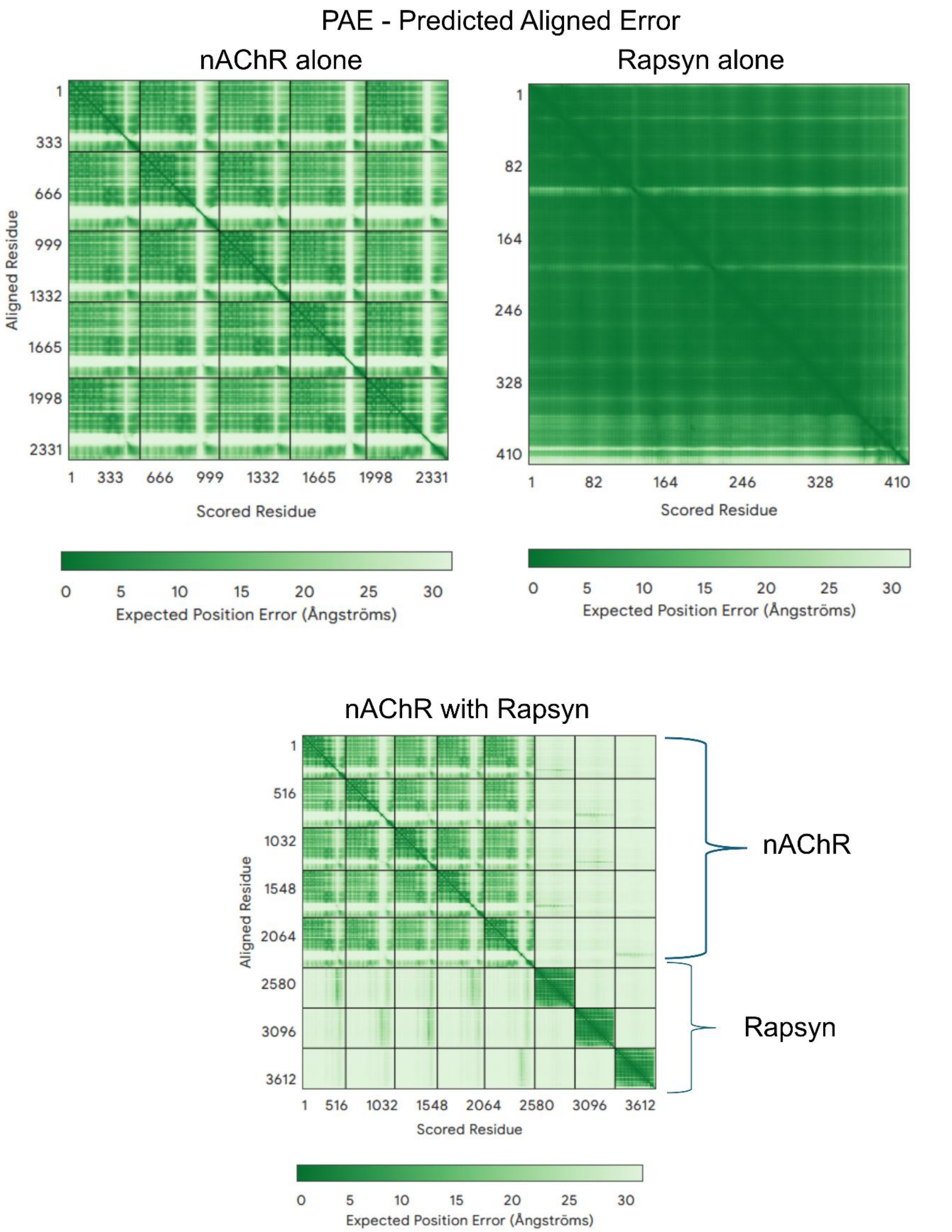


**Figure S8 Predicted aligned error** Representative AlphaFold3 PAE matrices for nAChR alone, rapsyn alone, and the 3-PTR-nAChR +3 rapsyn predictions.

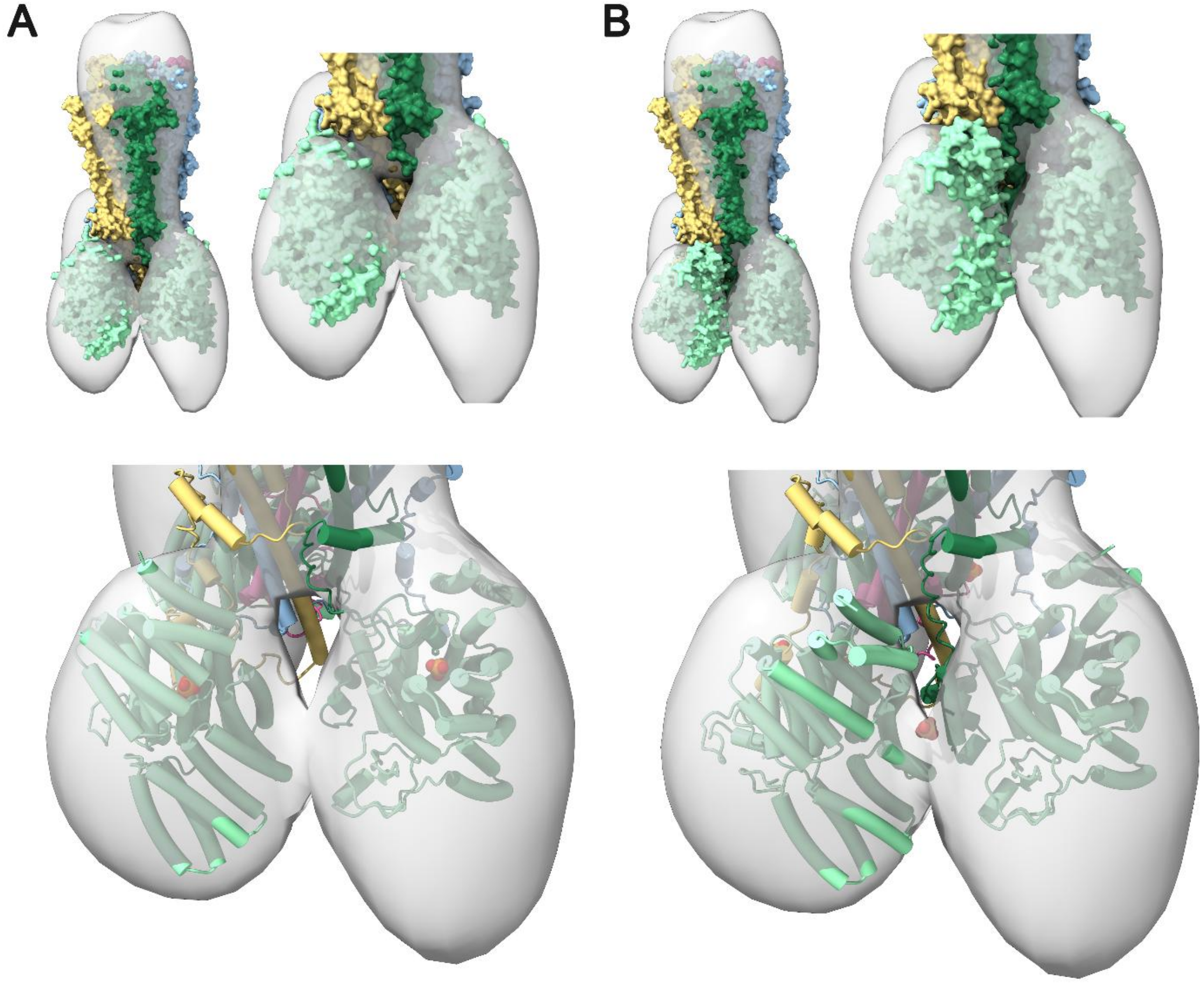


**Figure S9 AlphaFold3-predicted rapsyn–nAChR complexes fitted into the *Torpedo* cryo-ET density. (A)** Representative Alphafold3 model showing the predominant docking pose of rapsyn (teal) bound to the nAChR intracellular domain (β subunit) within the cryo-ET density from Zuber and Unwin (2013). In this orientation, the phosphate group (orange and red spheres) projects directly into rapsyn's positively charged binding pocket. This pose creates a gap between the rapsyn molecules, consistent with that seen in the map. This pose was seen in 37/45 rapsyn-nAChR interaction predictions. **(B)** Alternative predicted pose in which the phosphorylated tyrosine residue lies outside the binding pocket. This pose orients the two rapsyn molecules in closer proximity than the map suggests, filling the gap in the map.

Table S1 Cryo-EM data collection, refinement and validation statistics.

| | ICD Model | TightM4 | TiltM4 |
|---|---|---|---|
| **Data collection & processing** | | | |
| Microscope | | Glacios-IBS | |
| Magnification | | 36,000 | |
| Voltage (kV) | | 200 | |
| Frames (Exposure time in s) | | 40 (4) | |
| Total movies (no.) | | 9,810 | |
| Electron dose (e/Å$^2$) | | 42 | |
| Collection mode | | Counting | |
| Effective pixel size (Å) | | 1.1 | |
| Initial particle images (no.) | | 4,133,568 | |
| Final particle images (no.) | 139,486 | 48,744 | 90,742 |
| Symmetry imposed | C1 | C1 | C1 |
| Map resolution (Å) | 3.04 | 3.16 | 3.09 |
| **Model refinement** | | | |
| Initial model used (PDB code) | 7QL5 | 7QL5 | 7QL5 |
| **Model composition** | | | |
| Non-hydrogen atoms | 17,270 | 17,265 | 17,264 |
| Protein residues | 2,073 | 2,072 | 2,072 |
| Carbohydrates | 37 | 37 | 37 |
| Lipids | 0 | 0 | 0 |
| **B-factors** (min/max/mean Å$^2$) | | | |
| Protein | 108.8/254.32/150.7 | 113.2/276.46/159.42 | 114.66/274.29/158.84 |
| Ligand | 122.8/327.68/193.98 | 135.21/355.5/209.1 | 131.58/334.11/204.54 |
| **R.M.S. Deviations** | | | |
| Bond length (Å) | 0.002 (0) | 0.002 (0) | 0.002 (0) |
| Bond angles ($^0$) | 0.500 (0) | 0.500 (0) | 0.582 (0) |
| **Validation** | | | |
| MolProbity score | 1.54 | 1.64 | 1.59 |
| Clashscore | 4.46 | 5.62 | 5.65 |
| Rotamer outliers (%) | 0.00 | 0.00 | 0.00 |
| **Ramachandran plot** (%) | | | |
| Favored | 95.47 | 95.22 | 95.86 |
| Allowed | 4.53 | 4.78 | 4.14 |
| Outliers | 0.00 | 0.00 | 0.00 |
| Model Resolution | 3.6 | 3.7 | 3.7 |

Table S2. Quality scores of AlphaFold3 models of the nAChR without, and with increasing numbers of, rapsyn.

| Rapsyn | nAChR | PTMs present | ipTM | pTM |
|---|---|---|---|---|
| 0 | 1 | No | 0.71 | 0.71 |
| 1 | 1 | No | 0.61 | 0.64 |
| 2 | 1 | No | 0.56 | 0.59 |
| 3 | 1 | No | 0.50 | 0.53 |
| 4 | 1 | No | 0.49 | 0.52 |
| 1 | 1 | Yes | 0.60 | 0.63 |
| 2 | 1 | Yes | 0.57 | 0.60 |
| 3 | 1 | Yes | 0.50 | 0.53 |
| 4 | 1 | Yes | 0.49 | 0.52 |

Table S3. Residues added beyond the model 7QL5

| Subunit | Total additional residues | MX region | MA region |
|---|---|---|---|
| $\alpha_\gamma$ | 14 | A332 S333 K334 E335 | T364 P365 L366 I367 K368 N369 P370 D371 V372 K373 |
| β | 15 | P335 V336 T337 T338 | P394 V395 T396 L397 P398 Q399 D400 L401 K402 E403 A404 |
| δ | 10 | S341 R342 A343 D344 | D413 Q414 L415 H416 D417 E418 |
| $\alpha_\delta$ | 23 | M324 F325 F326 S327 T328 M329 K330 R331 | Q363 T364 P365 L366 I367 K368 N369 P370 D371 V372 K373 S374 A375 I376 E377 |
| γ | 14 | Q336 | D405 L406 A407 N408 F409 A410 P411 E412 I413 K414 S415 C416 V417 |
